# Geometric constraints within tripeptides and the existence of tripeptide reconstructions

**DOI:** 10.1101/2022.06.21.497005

**Authors:** Timothée O’Donnell, Viraj Agashe, Frédéric Cazals

## Abstract

Designing movesets providing high quality protein conformations remains a hard problem, especially when it comes to deform a long protein backbone segment, and a key building block to do so is the so-called tripeptide loop closure (TLC). Consider a tripeptide whose first and last segments (*N*_1_*C*_*α*;1_ and *C*_*α*;3_*C*_3_) are fixed, and so are all internal coordinates except the six {(*ϕ*, *ψ*)}_*i*=1,2,3_ dihedral angles associated to the three *C_α_* carbons. Under these conditions, the TLC algorithm provides all possible values for these six dihedral angles–there exists at most 16 solutions. TLC moves atoms up to ~ 5Å in one step and retains low energy conformations, whence its pivotal role to design move sets sampling protein loop conformations.

In this work, we relax the previous constraints, allowing the last segment (*C*_*α*;3_*C*_3_) to freely move in 3D space–or equivalently in a 5D configuration space. We exhibit necessary geometric constraints in this 5D space for TLC to admit solutions. Our analysis provides key insights on the geometry of solutions for TLC. Most importantly, when using TLC to sample loop conformations based on *m* consecutive tripeptides along a protein backbone, we obtain an exponential gain in the volume of the 5*m*-dimensional configuration space to be explored.

## 1 Introduction

### 1.1 Molecular dynamics and the Tripeptide Loop Closure

Biomolecular motions are inherently multi-scale, spanning ~ 15 and ~ 4 orders of magnitude in time and amplitude respectively [1]. Despite intensive efforts over the past fifty years or so, developing methods able to exploit this multi-scale structure has remained elusive. Two broad classes of methods have been developed. The first one relies on Newton’s equations, whose numerical solution uses time steps of the order of femto-seconds [2]. Alas, unless massive simulations are used [3] such tiny time steps are prohibitive for simulations with large systems, or systems undergoing large amplitude conformational changes. The second one, encompassing Monte Carlo based methods and basin-hopping like methods [2, 4], require *movesets* to propose novel conformations, which are accepted or not to further the simulation. While such methods are appealing since arbitrarily large spatial steps may be used, a shear difficulty is to avoid steric clashes and retain low (potential) energy conformations. This latter constraint is a strong incentive to work in internal coordinates, favoring dihedral angles which are *softer* coordinates than bond lengths and valence angles. In designing movesets, one classically distinguishes side chains and the protein backbone. To deal with side chains, various rotamer libraries have been used – see [5, 6, 7] and references therein. Dealing with the backbone is more challenging due to closure constraints arising when dealing with a loop squeezed in-between two *anchor* points.

A key building block in this context is the celebrated Tripeptide Loop Closure (TLC) algorithm [8, 9, 10]. In TLC, one consider three consecutive amino acids, with two types of constraints. The first one stipulates that the segments (also called *anchors* in the sequel) *N*_1_*C*_*α*;1_*C*_1_ and *N*_2_*C*_*α*;2_*C*_2_ are fixed in a given reference frame. The second one imposes that all internal coordinates are fixed, except the six rotatable bonds / dihedral angles {(*ϕ_i_*, *ψ_i_*)}_*i*=1,2,3_ found before / after the three *C_α_* carbons. Remarkably, TLC admits at most 16 solutions. This property sorts of discretizes the search space when running a simulation, in a manner akin to rotameric degrees of freedom for side chains.

Using TLC in the context of backbone simulations raises a difficulty, though. To see which, consider *m*(> 1) consecutive tripeptides. To use TLC on each tripeptide and mix their solutions if any, one needs to fix the position of all anchors. For a given tripeptide, this raises a novel question which is to study the existence of solutions to TLC when the second anchors moves relatively to the first one. This is precisely the problem addressed in this work, as specified now.

### 1.2 Contributions

We consider a tripeptide in which all internal coordinates (but the 6 dihedral angles) hold canonical values. Assuming that the two segments *N*_1_*C*_*α*;1_ and *C*_*α*;3_*C*_3_ are free to move with respect to one another, we aim at finding necessary conditions on these two segments for TLC to admit solutions. A tripeptide yielding solutions is termed *embeddable*. As we can assume without loss of generality that the first segment is fixed in a reference frame, this problem is posed in a five dimensional configuration space: the position of *C*_*α*;3_ enjoys 3 Cartesian coordinates, and that of *C*_3_ two spherical coordinates w.r.t. *C*_*α*;3_ (Fig. 1). The question becomes to find out which positions of *C*_*α*;3_ and *C*_3_ yielding embeddable tripeptides. To answer this question, our contributions are organized as follows:

- In Section 2, we present background material from [9].
- In Section 3, we derive *C_α_* valence constraints at each *C_α;i_* carbon to guarantee that the valence angle *θ_i_* at this *C_α_* is preserved. These constraints involve two angles denoted *σ*_*i*–1_ and *τ_i_*.
- In Sec. 4, we exploit a constraint associated with each *C_α;i_C*_*α;i*+1_ edge, to derive a deep validity constraint of iterative depth on all *σ*_*i*–1_ and *τ_i_* angles. This constraint is thus a necessary condition for the whole tripeptide
- Section 5 provides illustrations of our constructions, showing the sharpness of our constraints in the aforementioned five dimensional space.

**Figure 1:**
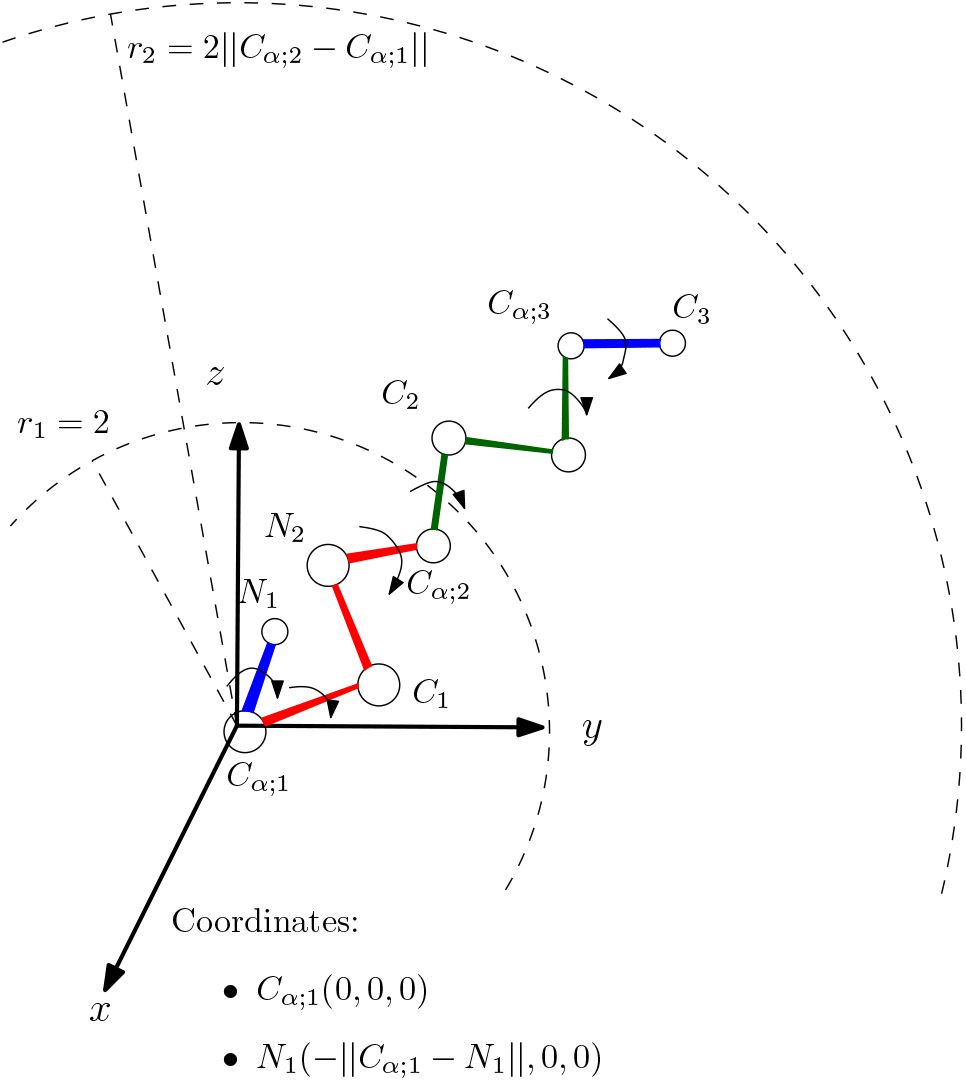
Reference frame for tripeptide embeddings. We consider a tripeptide whose internal coordinates are fixed, except the six {(*ϕ, ψ*)} dihedral angles associated with the three *C_α_* carbons. We assume that the segment *N*_1_*C*_*α*;1_ (first red line line segment) is fixed *i.e. C*_*α*;1_ is placed at the origin, and *N*_1_ is placed at (–||*N*_1_ – *C*_*α*;1_||, 0,0). We then aim at characterizing necessary conditions on the position of the last segment *i.e. C*_*α*;3_*C*_3_ for the Tripeptide Loop Closure (TLC) algorithm to hold solutions.

These contributions hinge on several interval types for the various angles involved in TLC (Fig. 2): IVI for Initial Validity Intervals, RVI for Rotated Validity Intervals, DVI for Depth-n/Deep Validity Intervals, RDVI for Restricted Deep Validity Intervals. In a sense, this work aims to understand the geometry of solutions of TLC in terms of necessary conditions on the six dihedral angles involved, expressed using these interval types.

**Figure 2:**
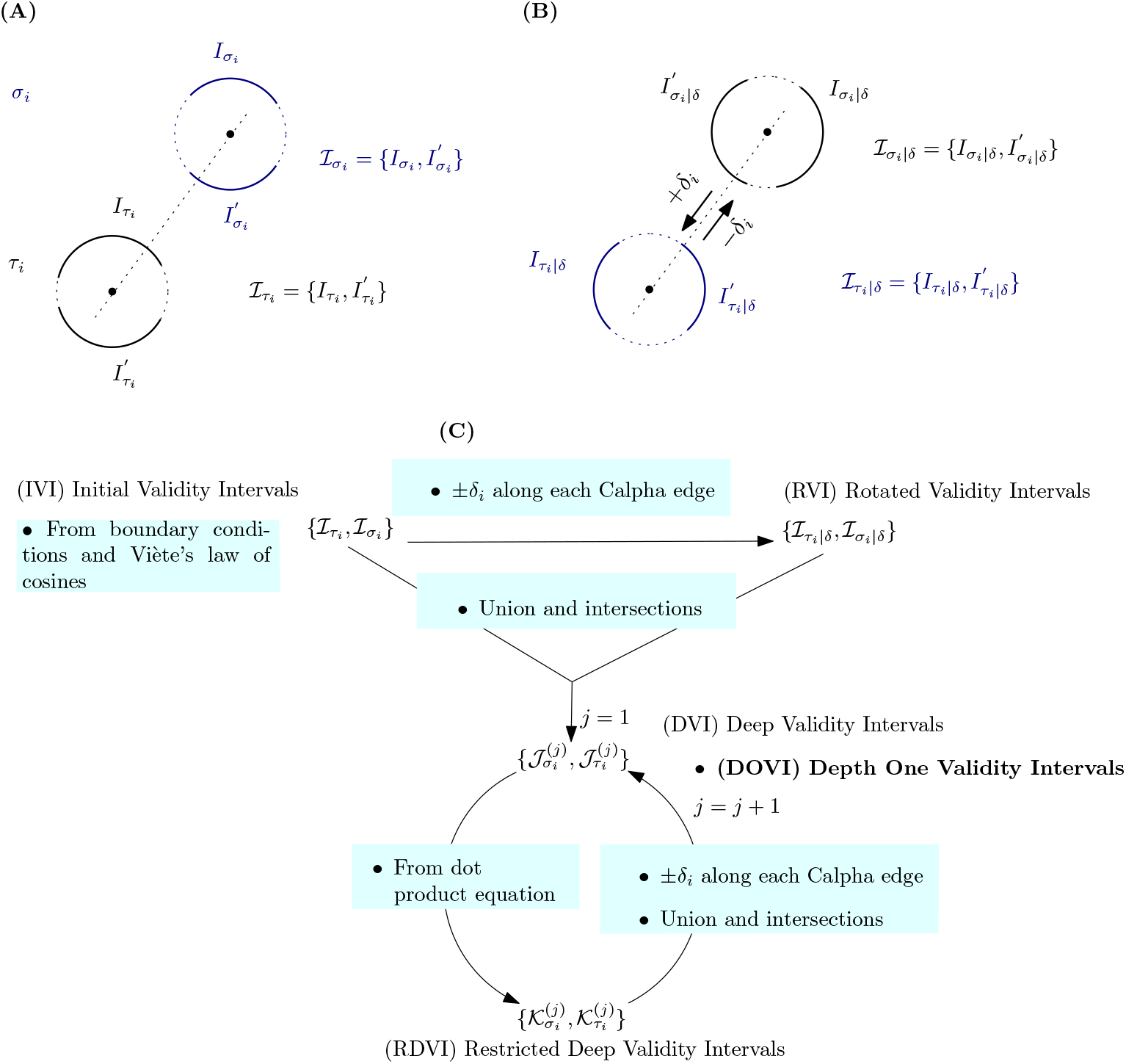
Validity interval types and their relationships. **(A) (IVI) Initial Validity Intervals.** See Def. 2. **(B) (TVI) Rotated validity intervals.** See Def. 3. Obtained from the initial validity intervals (**(A)**). **(C) Depth** *j***/Deep Validity Intervals and their restrictions.** From 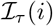 and 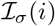 we obtain 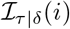 and 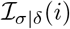. From all of those we obtain intersections constituting 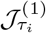 and 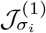. This *depth one validity interval* set can be refined to depth *n* iteratively (Def. 4, Algo 1).

## 2 Background on the Tripeptide Loop Closure

In this section, we review in detail the solution to TLC from [9].

### 2.1 Rotations and constraints

We consider the tripeptide (*N*_1_*C*_*α*;1_*C*_1_)(*N*_2_*C*_*α*;2_*C*_2_)(*N*_3_*C*_*α*;3_*C*_3_).

#### Geometry of the *C_α_* triangle of the tripeptide

Consider first the following four consecutive atoms *C*_*α*;1_*C*_1_*N*_2_*C*_*α*;2_ along the backbone, with *C*_1_*N*_2_ the peptide bond. Since the *ω*_1_ angle of the peptide bond is fixed, the distance ||*C*_*α*;1_ *C*_*α*;2_|| is constant (Fig. 3(A)). This observation holds for the other edge *C*_*α*;2_*C*_*α*;3_. Consequently, the atom *C*_*α*;2_ is restrained to the circle defined by the intersection of two spheres: *S*_1_(*C*_*α*;1_, ||*C*_*α*;1_*C*_*α*;2_||) and *S*_2_(*C*_*α*;3_, ||*C*_*α*;2_*C*_*α*;3_||).

**Figure 3:**
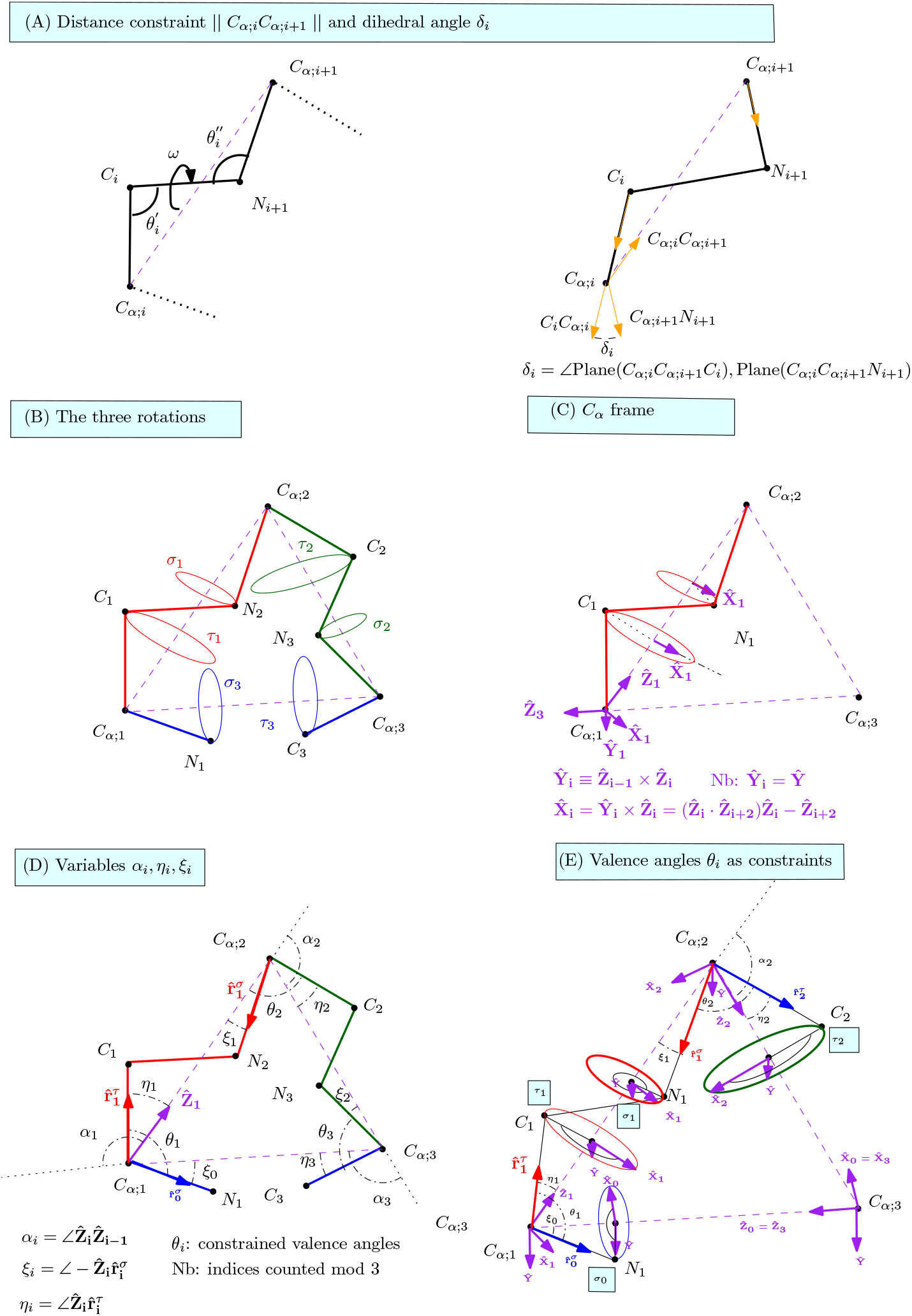
Tripeptide Loop Closure: main steps of the construction. Adapted from [9]. **(A)** Peptide bond linking two consecutive amino acids, and distance constraint induced on the line segment *C_α;i_C*_*α;i*+1_. The dihedral angle *δ_i_* is defined by the three vectors 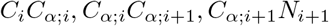. **(B)** The three rotations associated with the segments *C*_*α*;1_*C*_*α*;2_, *C*_*α*;2_*C*_*α*;3_ and *C*_*α*;3_*C*_*α*;1_. The rotation angles *τ_i_* (resp. *σ_i_*) concern atoms *C_i_* (resp. *N_i_*). But *τ_i_* and *σ_i_* satisfy *σ_i_* = *τ_i_* + *δ_i_*. **(C)** Construction of the local orthonormal frame associated with *C_α;i_* i.e. the *C_α_* frame. **(D)** Introducing the variables *α_i_, η_i_, ξ_i_*. **(E)** Modeling the constraint on valence angles at *C_α;i_* carbons.

Finally, consider the base of this triangle. By hypothesis, atoms *N*_1_, *C*_*α*;1_, *C*_*α*;3_, *C*_3_ are fixed, so that the length of the base is fixed.

The geometry of the triangle *C*_*α*;1_*C*_*α*;2_*C*_*α*;3_ is therefore fixed. However, one has one rotating rigid body attached to each of its edges (Fig. 3(B)):

- The movement of the atom *C*_1_ (resp. *N*_2_) can be modeled by a rotation of angle *τ*_1_ (resp. *σ*_1_) about the axis *C*_*α*;1_*C*_*α*;2_;
- The movement of the atom *C*_2_ (resp. *N*_3_) can be modeled by a rotation of angle *τ*_2_ (resp. *σ*_2_) about the axis *C*_*α*;2_*C*_*α*;3_;
- The movement of atoms *N*_1_*C*_*α*;1_*C*_*α*;3_*C*_3_ can be modeled by a rotation of angle *τ*_3_ about the axis *C*_*α*;1_*C*_*α*;3_.

##### Remark 1

*Using the dihedral angle δ_i_ defined by the four atoms C_α;i_, C_i_, N*_*i*+1_, *C*_*α;i*+1_ (*Fig. 3(B)) one has the relationship*:

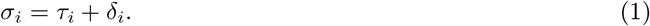

#### Local *C_α_* frames

The TLC from [9] defines one local frame to handle the rotation angles *τ_i_* and *σ_i_*. Using these frames and Eq. (1), TLC reduces the problem to three variables and three constraints:

- The three variables: the angles *τ_i_*∀*i* ∈ {1, 2, 3} which can be rotated.
- The three constraints: the valence angles *θ_i_*∀*i* ∈ {1, 2, 3} each at the corresponding *C_α;i_*, which which must be kept constant.

It is the coupling introduced by the *θ_i_* angles onto the rotation angles *τ_i_* that yields a degree 16 polynomial [9, 10].

We have recalled above that in TLC, the atoms *N*_1_, *C*_*α*;1_, *C*_*α*;3_, *C*_3_ are fixed. But in using the rotation angles {(*τ_i_, σ_i_*)}, the segments *N*_1_*C*_*α*;1_ and *C*_*α*;3_*C*_3_ are moving. Therefore, once the atomic positions have been obtained using the local frames, all atoms are rotated such that these four are back into their original positions in the main frame.

##### Remark 2

*The valence angles around C_i_ and N_i_ atoms do not play a role, since they are fixed and internal to the two segments being rotated*.

### 2.2 Local coordinate system at *C_α;i_*

Three **unit** vectors 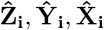 are defined to form an orthonormal coordinate system (Fig. 3(C)). These vectors are:

- (Unit vector aligned with one side of the triangle) 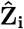 is the unit vector anchored at *C_α;i_* aligned with the edge of the *C_α_* triangle. Note that *C*_*α*;3_ points to *C*_*α*;1_.
- (Unit vector perpendicular to the plane of the triangle) 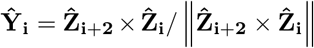. Intuitively, note that 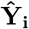 is obtained by taking the cross product between two of the unit vectors just mentioned: that *entering* the *C_α_* carbon and that *exiting* it. (Nb: the vector 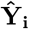 does not depend on the index *i*, which can be dropped.)
- (Reference unit vector to define the rotation angles) 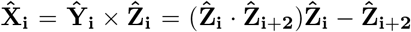. (For the latter, we use the double cross product formula *u* × (*v* × *w*) = (*u.w*)*v* – (*u.v*)*w*.)

Using the local frames, one also defines the angles *α_i_, η_i_* and *ξ_i_* as follows (Fig. 3(D)):

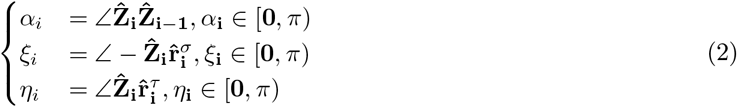

These variables will be used to handle the conservation of the valence angle at *C_α;i_*, which requires considering three atoms: *N*_*i*–1_, *C_α;i_*, and *C_i_*.

#### Remark 3

*Indices i* ∈ {1, 2, 3} *are counted modulo 3; that is, i* – 1 *is equivalent to i* + 2: *e.g*., 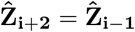.

#### Remark 4

*The following equalities are used throughout all calculations*^1^:

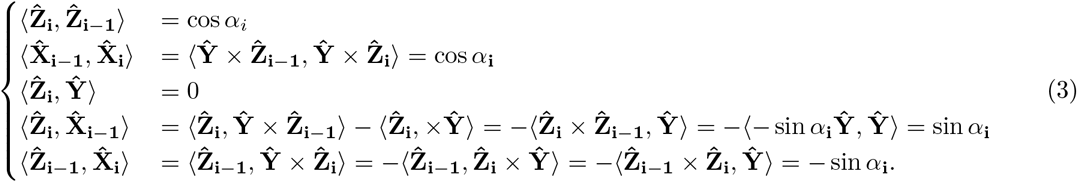

### 2.3 Rotations of *N_i_* and *C_i_*

Consider 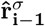 the *C_α;i_N_i_* unit vector and 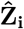 the *C_α;i_C_i_* unit vector. The rotations of *C_i_* and *N_i_* are described as follows (Fig. 3(C,D)):

- atom *N_i_*, angle *σ*_*i*–1_: rotation of 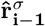 about 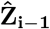
- atom *C_i_*, angle *τ_i_*: rotation of 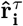 about 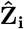

The vectors 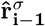 and 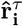 are easily obtained using the local frames (Fig. 3(E)):

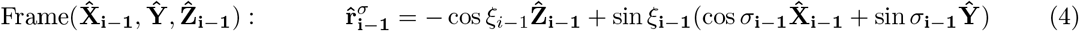

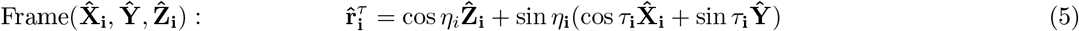

Using the previous two equations and the equalities from Eq. (3), one obtains the dot product between 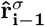 and 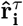:

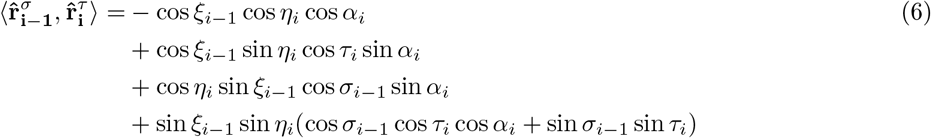

The conservation of the valence angle *θ_i_* imposes the following *valence angle constraint*: – see also Fig. S1:

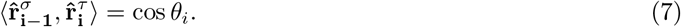

In TLC only the *τ_i_* and *σ*_*i*–1_ terms vary, the value of this dot product can thus be represented depending on those two angles.

#### Remark 5

*It should be noticed that the angles of Eq. (2) and the* {*σ,τ*} *angles serve complementary purposes: the angles ξ, η define cones for the rotating atoms N and C, while the angles* {*σ, τ*} *position the rotating atoms onto a circle orthogonal to the axis of each such cone*.

### 2.4 Internal coordinates and the 5D configuration space

As mentioned in Sec. 1.2, we study TLC when the position of the first two and last two atoms are given, and all internal coordinates (but *ϕ*, *ψ* angles) are fixed. (Note that fixing the position of the segment *C*_*α*;3_*C*_3_ requires 5 parameters, whence a 5D configuration space.) With these two pieces of informations, we compute all angles which define a unique TLC problem. To obtain the 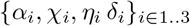 values which are sufficient to solve TLC, we proceed as follows. We first note that *η*_3_, *ξ*_3_ and *δ*_3_ are directly obtained from the positions of the anchors *N*_1_*C*_*α*;1_ and *C*_*α*;3_*C*_3_. Fixed internal coordinates yield the geometry of the *C_α_* triangle, whence the *α_i_* values. Finally, using the geometry of the *C_α_* triangle and that of the peptide bodies *C_α_CNC_α_*, one obtains *η_i_, ξ_i_, δ_i_* for *i* ∈ 1..2.

## 3 *C_α_* valence angle constraints

In this section, we study constraints on *σ*_*i*–1_ and *τ_i_* so as to guarantee that the valence angle *θ_i_* is preserved. We first present derivation of intervals for *σ*_*i*–1_ and *τ_i_* (Sec. 3.1), and proceed with the no-solution case, introducing a necessary condition for solutions to exist (Sec. 3.2).

### 3.1 Initial validity intervals for *σ*_*i*–1_ and *τ_i_*

We first derive initial validity intervals, using boundary conditions at each *C_α_* carbon. The construction for *σ*_*i*–1_ uses two quantities *S*^−^ and *S*^+^, from which both existence conditions and the angular values are derived (Fig. 4). The analysis proceeds mutatis mutandis for *τ_i_*.

**Figure 4:**
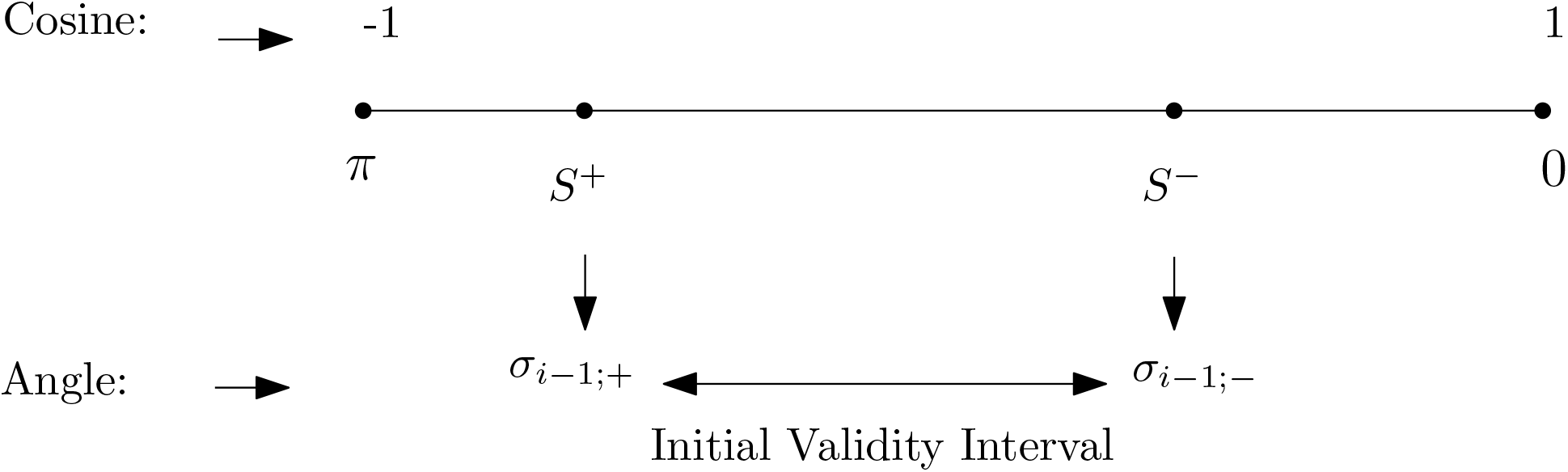
Conditions to define the extreme angles involved in the initial validity intervals: the case of *σ*_*i*–1_.

#### 3.1.1 Angle *σ*_*i*–1_

We wish to define a *validity interval* for *σ*_*i*–1_, namely *I*_*σ*_*i*–1__ = [*σ*_*i*–1;–_, *σ*_*i*–1;+_] ⊆ [0, *π*].

##### Basic equation

We first wish to restrict the values of *σ*_*i*–1_ for which *θ_i_* can be preserved, and use to this end the reference vector 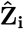. Using the expression of 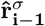 in the local frame of *C*_*α;i*–1_ (Eq. (4)), we get:

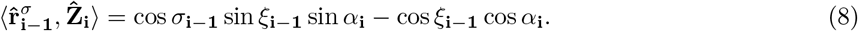

Independently from the angle *σ*_*i*–1_, elementary spherical trigonometry based on Viète’s law of cosines makes it possible to bound the angle 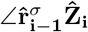 as follows (Sec. S1.2, Fig. S2):

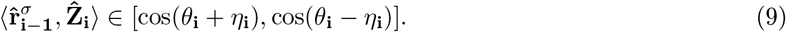

Exploiting the bounds of the interval yields two extreme values for the angle *σ*, denoted *σ*_*i*–1;–_ and *σ*_*i*–1;+_.

##### Angle *σ*_*i*–1;–_

The first limit case reads as:

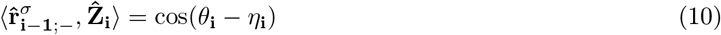

from which we obtain

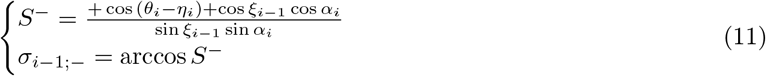

When *S*^−^ → 1^−^ by properties of arccos, we have *σ*_*i*–1;–_ → 0^+^ (Fig. 4). Therefore, when

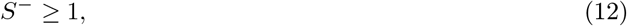

we set *σ*_*i*–1;–_ = 0, so that any value *σ*_*i*–1_ ≤ *σ*_*i*–1;+_ is valid.

##### Angle *σ*_*i*–1;+_

The second limit case reads as:

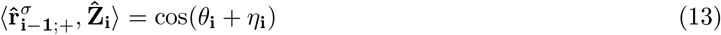

from which we obtain

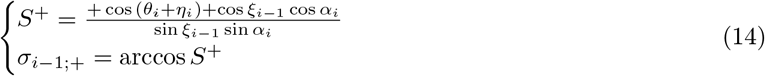

When *S*^+^ → –1^+^, by properties of arccos, we have *σ*_*i*–1;+_ → π^−^. Therefore, when

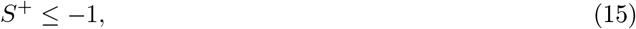

we set *σ*_*i*–1;+_ = π, so that any value *σ*_*i*–1_ ≥ *σ*_*i*–1;–_ is valid.

##### Illustration

When considering the dot product 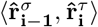 as a function of the two variables *τ_i_* and *σ*_*i*–1_, the angles *σ*_*i*–1;–_ and *σ*_*i*–1;+_ correspond to planes orthogonal to the *σ*_*i*–1_ axis (Fig. 5(B,C)).

**Figure 5:**
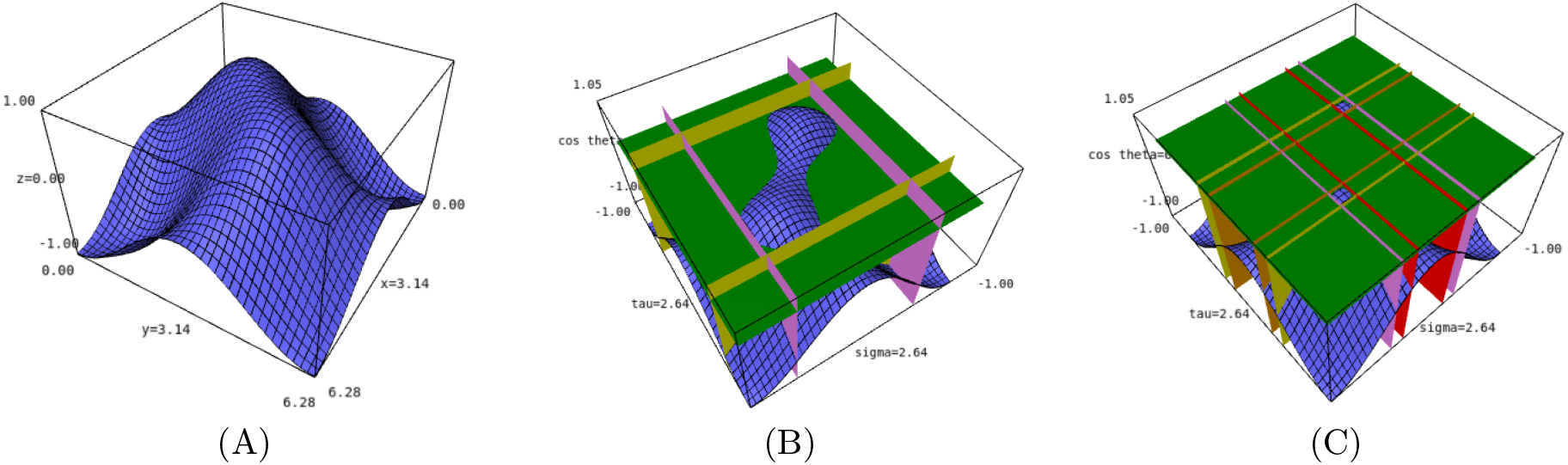
Example dot product surface and extreme angles 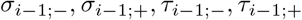. TLC problem for the values *α_i_* = 100, *χ*_*i*–1_ = 50, *η_i_* = 50 **(A)** Whole surface **(B)** With horizontal plane cos *θ_i_* = cos 9°. Note the four vertical planes corresponding to the extreme angles. In this case, the intersection between the surface and the plane consists of a plane curve with two connected components. One component is enclosed by the four vertical planes. **(C)** With horizontal plane cos *θ_i_* = cos 35°. In this case, the intersection between the surface and the plane consists of a plane curve with one connected component.

#### 3.1.2 Angle *τ_i_*

We also wish to set a *validity interval* for *τ_i_*, that is 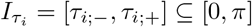.

##### Basic equation

We proceed mutatis mutandis for the vector 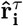, using vector 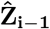 as landmark. Using the expression of 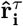 in the local frame of *C_α;i_* (Eq. (5)), one obtains:

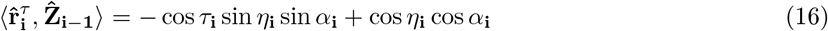

Viète’s law of cosines now yields the following for the angle 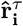:

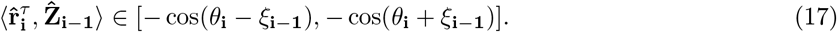

Combining the previous two equations yields up to two solutions, namely *τ*_*i*;–_ and *τ*_*i*;+_.

##### Angle *τ*_*i*;–_

The first limit case reads as:

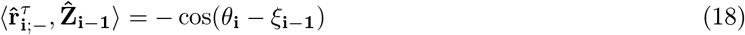

from which we obtain

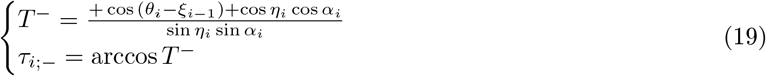

When *T*^−^ → 1^−^, by properties of arccos, we have *τ*_*i*;–_ → 0^+^. Therefore, when

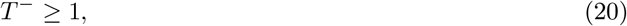

we set *τ*_*i*;–_ = 0, so that any value *τ_i_* ≤ *τ*_*i*;+_ is valid.

##### Angle *τ*_*i*;+_

The second limit case reads as:

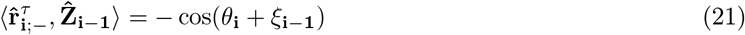

from which we obtain

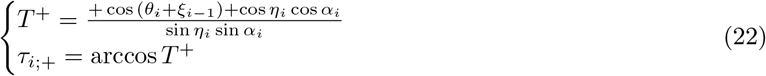

When *T*^+^ → –1^+^, by properties of arccos, we have 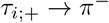. Therefore, when

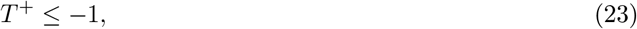

we set *τ*_*i*;+_ = *π*, so that any value of *τ_i_* ≥ *τ*_*i*;–_ is valid.

##### Illustration

When considering the dot product 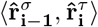 as a function of the two variables *τ_i_* and *σ*_*i*–1_, the angles *τ*_*i*–1;–_ and *τ*_*i*–1;+_ correspond to planes orthogonal to the *τ_i_* axis (Fig. 5,(B),(C)).

### 3.2 Necessary conditions for *σ*_*i*–1_ and *τ_i_*

In deriving the lower and upper bounds of the initial validity intervals for *σ* and *τ*, we already processed four limit cases for the dot products (Eqs. (12), (15), (20), (23)). The remaining four yield the following:

#### Definition. 1

*The C*_*α*_ valence constraints *are the necessary validity conditions defined by*:

- Angle *σ*_*i*–1;–_: *the condition σ*_*i*–1;–_ < *σ*_*i*–1;+_ *requires*

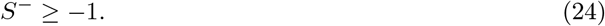
- *Angle σ*_*i*–1;+_: *the condition σ*_*i*–1;–_ < *σ*_*i*–1;+_ *requires*

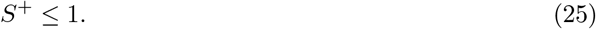
- *Angle τ*_*i*;–_: *the condition τ*_*i*;–_ < *τ*_*i*;+_ *requires*

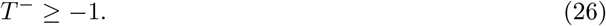
- *Angle τ*_*i*;+_: *the condition τ*_*i*;–_ < *τ*_*i*;+_ *requires*

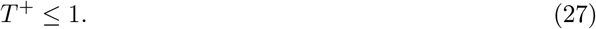

*For the constraint to be verified all these conditions must be valid for three pairs* {(*σ*_*i*-1_, *τ_i_*)} *associated with the three C*_*α*_ *carbons*.

Summarizing, when the previous equations are not verified, no initial validity interval can be defined for *σ*_*i*–1_ and/or *τ_i_* (Fig. 6(E) Fig. 4).

**Figure 6:**
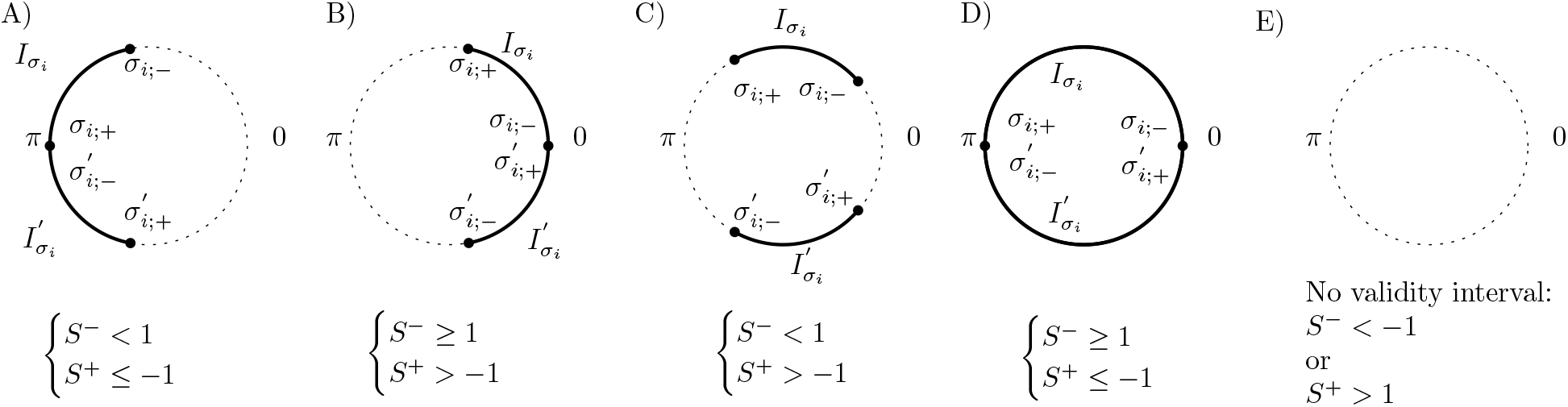
The four possible types of *initial/rotated* validity interval types for the angle *σ*. The quantities of interest are defined in Eqs. (12) (15), (24), (25). Cases (A) to (D) stand for situations where *σ*_*i*;–_ and/or *σ*_*i*;+_ can be defined. In case (E), no validity interval can be defined.

### 3.3 Symmetry around the *C_α_* triangular plane, and *C_α_* valence constraints

The previous angles are defined in [0, π]. Due to the symmetry of the tripeptide with respect to the *C_α_* plane, these angles have counterparts in [π, 2π]. We therefore define the following symmetric intervals:

- The *symmetric validity interval* for *σ*_*i*–1_ is defined by

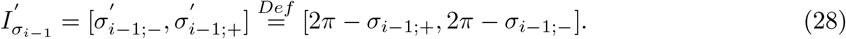
- The *symmetric validity interval* for *τ_i_* is defined by

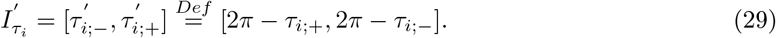

Using these, we can finally specify the valid intervals for the angles *σ*_*i*–1_ and *τ_i_* must belong to:

#### Definition. 2

*The* initial validity interval set *for σ*_*i*–1_ *is defined by*:

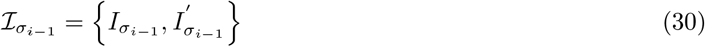

*Likewise, the initial validity interval set for τ_i_ is defined by*:

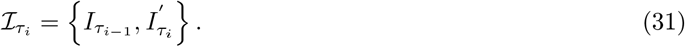

For *σ*_*i*–1_, as long as the conditions of Eqs. (24) and (25) are satisfied, we have:

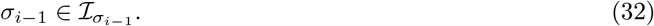

For *τ_i_*, as long as the conditions of Eqs. (26) and (27) are satisfied, we have:

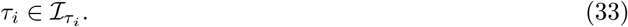

Combining the previous conditions yields the complete case analysis (Fig. 6).

## 4 Deep validity constraints associated with the *C_α_* triangle

### 4.1 Exploiting the coherence along a *C_α;i_C*_*α;i*+1_ edge

The constraints presented in the previous section focus on the three *C_α_* carbons independently. On the other hand, for a given tripeptide, the angles *τ_i_* and *σ_i_* and the dihedral angle *δ_i_* defined by the four atoms *C_*α;i*_C_i_N*_*i*+1_*C*_*α*;*i*+1_ satisfy *σ_i_* = *τ_i_* + *δ_i_* (Eq. (1)).

Given an interval of values for *τ_i_*, the previous formula can be used to infer a projected interval for *σ_i_*, and vice-versa (Fig. 2(B)). Whence the following definitions which exploit the previous formula along the two edges of the *C_α_* triangle incident on *C_α;i_*:

#### Definition. 3

*The* rotated validity interval set *for the angles *σ*_*i*–1_ and τ_i_ are defined by*:

- *for* 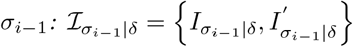 *with*:

– 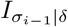: *interval for σ*_*i*–1_ *obtained by applying Eq. (1) to I*_*τ*_*i*–1__. (*Nb: uses the edge C_α;i_C*_*α;i*–1_ *of the C_α_ triangle*.)
– 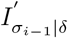: *interval for σ*_*i*–1_ *obtained by applying Eq. (1) to* 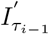. (*Nb: uses the edge C_α;i_C*_*α;i*–1_ *of the C_α_ triangle.*)
- *for* 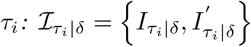 *with*:

– 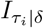: *interval for τ*_*i*_ *obtained by applying Eq. (1) to I*_*σ*_*i*__. (*Nb: uses the edge C_α;i_C*_*α;i*+1_ *of the C_α_ triangle*.)
– 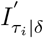: *interval for τ*_*i*_ *obtained by applying Eq. (1) to* 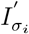. (*Nb: uses the edge C_α;i_C*_*α;i*+1_ *of the C_α_ triangle*.)

Summarizing, for each of the *σ*_*i*–1_ and *τ_i_* angles, we have obtained 4+4 intervals using the *θ_i_* angle constraint at *C_α;i_* (Eq. (6)):

- Four for the *σ*_*i*–1_ angle: 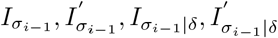
- Four for the *τ_i_* angle: 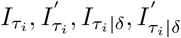

### 4.2 Deep validity Intervals: depth one

We are now in position to combine two pieces of information:

- The conditions on *σ*_*i*–1_ and *τ_i_* inherent to the conservation of the valence angles (Eq. (7)).
- The conditions exploiting rotated validity intervals, stemming from Eq. (1)

Since the previous intervals define necessary conditions, intersections between intervals for a given angle must be non empty. We therefore combine them as follows 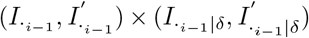, which yields *depth 1 validity intervals*:

#### Definition. 4

*The* depth 1 validity interval set 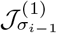 *for σ*_*i*–1_ *is*:

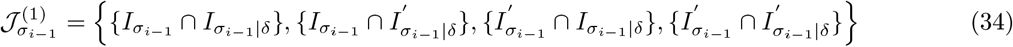

*Similarly, the depth 1 validity interval set* 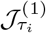 *for τ_i_*:

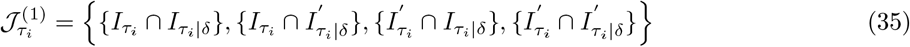

Note that each depth 1 validity interval has up to four connected components.

#### Definition. 5

*The* depth 1 validity constraint *for σ*_*i*–1_ reads as:

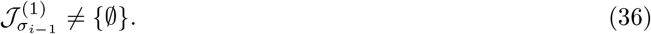

*The depth 1 validity constraint for τ_i_ reads as*:

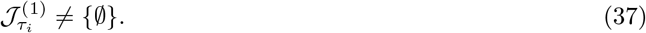

*The* depth 1 validity constraint *for the tripeptide is the conjunction of the constraints for the six σ and τ angles*.

As we shall see, this constraint is significantly more restrictive than the corresponding *C_α_* valence constraint (Def. 1, Fig. 7).

**Figure 7:**
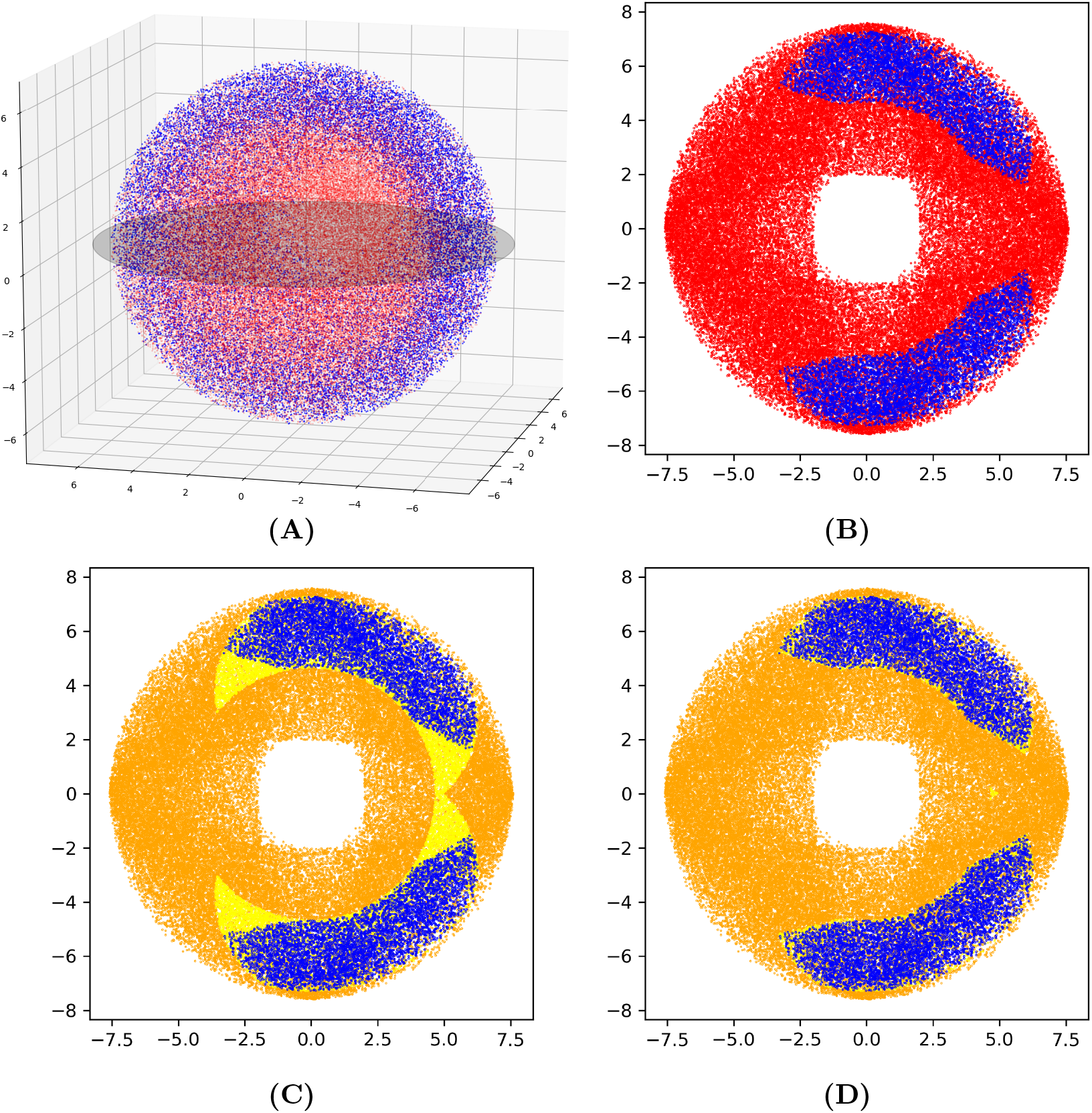
Embeddable tripeptides and necessary conditions: stringency of *C_α_* valence constraints (Def. 1) versus depth 1 validity constraints (Def. 5), illustrated on random instances projected into the reference frame of Fig. 1. (Nb: figures in 3D and 2D, while the configuration space is 5D.) **(A)** Blue (resp. red) points represent positions of *C*_*α*;3_ in instances when TLC yields at least one solution (resp. yields no solution). **(B)** A similar dataset generated uniformly on the of the sphere–gray equator in (A), color code as in (A). **(C)** *C_α_* **valence constraints.** The *C*_*α*;3_ positions are depicted using three colors: blue points as in (A,B); orange points: points failing the *C_α_ valence constraints;* yellow points: points satisfying the *C_α_ valence constraints*, but for which TLC admits no solution. **(D) *Depth 1* validity constraints.** Color code as in (C), using the depth 1 validity constraints instead of the *C_α_* valence constraints. Note the reduction of the yellow region.

The derivation of these validity constraints can be seen as a bootstrap process (Fig. 2). The initialization consists of computing the initial validity intervals, using the boundary conditions imposed by Equations (11) (14) (19) (22). The second step consists of exploiting the coherence along each *C_α_* edges, as imposed by Eq. (1).

It should also be noticed that transposing the initial validity intervals from *τ_i_* to *σ_i_* using Eq. (1) is equivalent to transposing from *σ_i_* to *τ_i_* using the same equation. Therefore the *depth 1 validity constraint* for *σ_i_* and the one for *τ_i_* are redundant.

#### Remark 6

*The previous intersections naturally depend on the two anchor positions and the fixed internal coordinates*.

### 4.3 Deep Validity Intervals: arbitrary depth

The qualifier *depth 1* used in the previous section indicates that the dual process consisting of specifying validity intervals at each *C_α_* and projecting along a *C_α_* edge can be repeated, moving from necessary conditions at depth *j* to necessary conditions at depth depth *j* + 1.

#### 4.3.1 Functions *σ*_*i*–1_(*τ_i_*) and *τ_i_*(*σ*_*i*–1_)

We first note that an interval for *σ*_*i*–1_ or *τ_i_* implies two intervals for the second angle – obtained by computing *σ*_*i*–1_ from *τ_i_*, or vice versa. To see how, note that the dot product equation (6) can be written as:

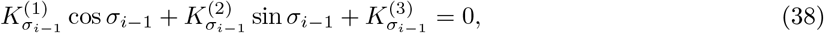

with

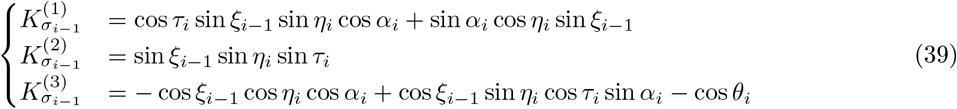

As explained in Sec. S1.4, for a given *τ_i_*, with atan2(*y*, *x*) = arg(*x* + *iy*), the two possible solutions are given by:

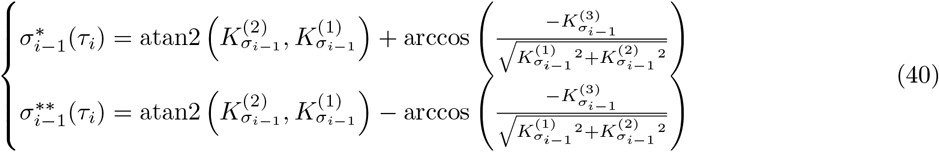

Therefore, given a value of *τ_i_*, we obtain two values of *σ*_*i*–1_ which validate the valence angle constraint by using the above equations and vice-versa.

Using the previous, we define:

##### Definition. 6

*For each interval* 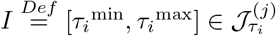, *consider the two intervals*

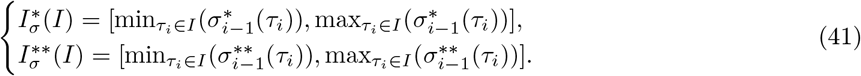

*The* depth *j* validity interval set *is defined by*:

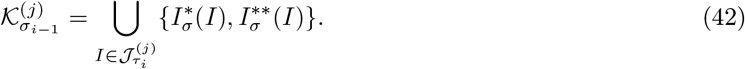

One proceeds mutatis mutandis to obtain the values and intervals for 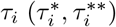 given *σ*_*i*–1_, as well as 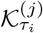.

At this point, let us recap on the two complementary routes used to study the angles *σ*_*i*–1_:

- the dot product of Eq. (6) and the conservation equality 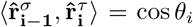 can be rewritten as Eq. (38), from which one derives two continuous functions *σ*_*i*–1_ as a function of *τ_i_*.
- the dot product 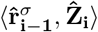 (Eq. (8)) yields tight upper and lower bounds on *σ*_*i*–1_ using Viète’s law of cosines – SI Sec. S1.2.

The bounds of the former functions match the latter values, which we summarize as follows:

##### Observation. 1

*The extrema of the functions* 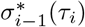 *and* 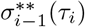 *are the bounds of the initial validity intervals, as specified in Eqs. (11) and (14)*.

Illustrations for these functions are provided in the Sec. S1.4.

#### 4.3.2 Iterated intervals

Given the depth 1 validity intervals for *σ*_*i*–1_, a set of intervals can be defined for *τ_i_*. A temporary set of intersections between this set and the depth 1 validity intervals for *τ_i_* can then be defined. Finally using intersections between temporary sets and rotated temporary sets (Eq. (1)) we can obtain *depth 2 validity intervals* (Fig. 2(C)). This iterative process is summarized in Algo. 1.

##### Algorithm 1 Computing depth *j* validity intervals

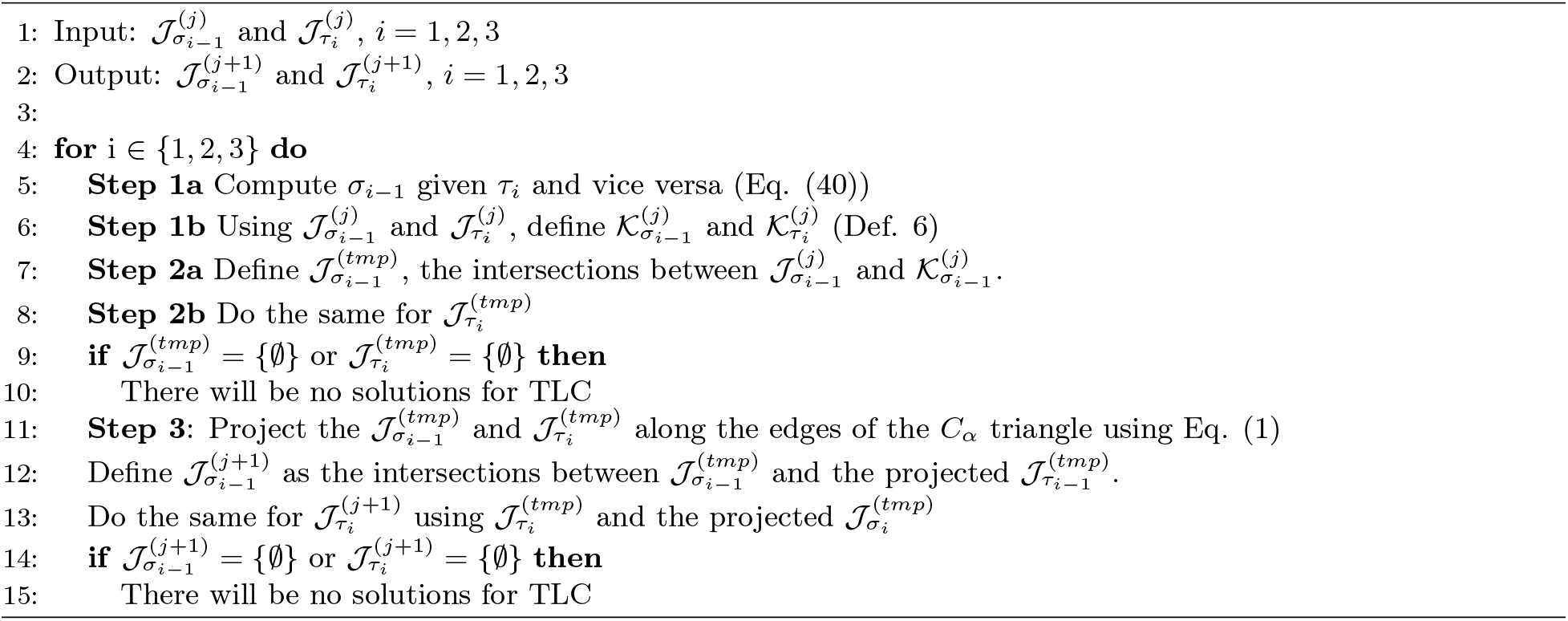

##### Maximum number of deep validity intervals (DVI)

###### Lemma. 1

*Let n_j_ be the maximum number of intervals at depth j in* 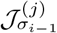 *or* 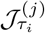. *This number satisfies*

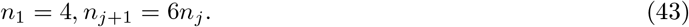

*Proof*. We first raise an observation for two sets of intervals 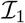 (of size *n*_1_) and 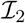 (of size *n*_2_) such that 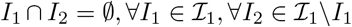. Assuming, without loss of generality, that *n*_2_ ≥ *n*_1_, the maximum number of intervals determined by the intersections between intervals in these sets is 2 × *n*_1_ + *n*_2_ – *n*_1_ for intervals on a circle. To build this worst-case, we stab two intervals in 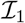 with an interval in 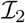, and squeeze the remaining *n*_2_ – *n*_1_ intervals from 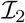 inside intervals of 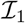.

To establish the lemma, we follow the steps of Algo. 1, starting with the set 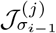. If needed, we redefine the intervals in this set so that they are disjoint – to meet the hypothesis on the two sets 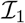 and 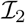 used in the observation above.

Considering the *n_j_* intervals making up 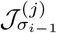 (or 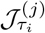), we now inspect the following steps of Algo. 1:

- **Step 1b**) projects 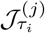 to yield 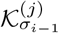 (Def. 6). By definition this yields 2*n_j_* restricted validity intervals.
- **Step 2a**) takes the intersection between 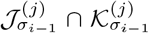, which the observation above, yields 2*n_j_* + (2*n_j_* – *n_j_*) = 3*n_j_* intervals.
- **Step 3**) takes the intersection between the 3*n_j_* intervals of the last step and the 3*n_j_* from the δ projected intervals. This yields a maximum of 6*n_j_* intervals.

## 5 *C_α_* valence constraint and depth 1 validity constraints: illustrations

### 5.1 Material: dataset of random instances

Our experiments use standard internal coordinates for bond lengths and valence angles [9]. The canonical values are available in our TLC implementation user manual (https://sbl.inria.fr/doc/Tripeptide_loop_closure-user-manual.html).

#### General dataset

We place our tripeptide in the reference frame using the first segment (Fig. 1). In this frame, we randomly generate *C*_*α*;3_ between two spheres around the origin. The inner radius *r*_1_ = 2Å is smaller than the smallest value ||*C*_*α*;3_ – *C*_*α*;1_| found in our exhaustive database of ~ 2.5 million tripeptides extracted from the PDB [11]. The outer radius *r*_2_ = 2 ||*C*_*α*;2_ – *C*_*α*;1_|| respects the triangle inequality based on the distances between *C_α_* carbons, and the canonical internal coordinates values. The position of *C*_*α*;2_ is generated uniformly in this volume. Atom *C*_3_ is generated on a sphere centered at *C*_*α*;3_, with a radius defined by the canonical bond length. The positions of *N*_1_*C*_*α*;1_ and *C*_*α*;3_*C*_3_ together with the canonical internal coordinate yield a TLC problem, which is fertile if embeddings/solutions are obtained. To each of those inputs correspond values for the input angles *α_i_, ξ_i_*, *η_i_* for each *C_α;i_*.

Over 100,000 instances, 24,076 fertile ones were observed.

#### Planar dataset

A second similar dataset of the same size with *C*_*α;i*+2_ being positioned uniformly between two circles using the same radii. *C*_*i*+2_ is then also generated on a sphere around *C*_*α;i*+2_.

### 5.2 Validity intervals

#### Signatures

To assess the diversity of situations faced at *C_α_* carbons, we introduce a signature based on the *σ* and *τ* interval types:

##### Definition. 7

*At a C_α;i_ carbon, consider the endpoints of the validity intervals, in this order σ*_*i*–1;–_, *σ*_*i*–1;+_, *τ*_*i*;–_, *τ*_*i*;+_. *The* signature *of a TLC problem is a string in* {*N,P,Z*}^4^ – *one letter for each each extreme angle, with the following convention*:

- *letter N for cos* (*endpoint*) < −1,
- *letter P for cos* (*endpoint*) > 1,
- *letter Z for* −1 < *cos*(*endpoint*) < 1.

Among the sample of embeddable anchor positions for TLC, only seven different signatures are found (Fig. S3). Not all combinations are possible as the the first and third letter cannot be N (Eq. (26)). Similarly the second and fourth cannot be P (Eq. (27)). If the first and second are ZZ (resp. third and fourth) then their counterpart will be PN. This illustrates that if the two endpoints are defined in]0, π[for *σ*_*i*–1_, then the whole circle is possible for *τ_i_* (and vice versa).

### 5.3 Necessary conditions and validity constraints

#### General dataset

Using the general dataset, plotting the positions of *C*_*α*;3_ associated with solutions yields a bell shaped distribution with a hole on top (Fig. 7(A)).

#### Planar dataset

Due to the symmetry around the *N*_1_*C*_*α*;1_ line, we consider the aforementioned planar data set (Fig. 7(B)). Consider a set of conformations, some fertile (TLC admits solutions), and some sterile. Fertile conformations naturally satisfy the necessary conditions defined by both the *C_α_ valence constraints* (Def. 1) and the depth 1 validity constraints (Def. 5) for all *σ* and *τ* angles. However, we would like these constraints to be as tight as possible, retaining as few sterile configurations as possible. To assess this, for each constraint, we color code sterile configurations (Fig. 7(B), red points) using two colors: orange for sterile but failing the necessary test, and yellow for sterile passing the necessary test. Ideally, we would like as few yellow points as possible. It clearly appears that the *depth 1 validity constraints* (Fig. 7(D)) are much tighter than the *C_α_ valence constraints* (Fig. 7(C)).

In terms of proportions, 24.08% of the points are instances where TLC yields solutions. In total 41.35% fulfill the *C_α_* valence constraints. Finally 28.12% fulfill the *depth one validity constraints*. They are included in the 41.35% and include the 24.08% just mentioned. The cases validating the depth one validity constraint therefore represent ~ 2/3 of the cases fulfilling the *θ_i_* equations. For the sake of clarity, it should be stressed that the gap between the 24.08% of instances with solutions, and the 28.12% fulfilling the depth one validity constraints is not clearly visible using our projections in 3D and 2D (Fig. 7) since the whole configuration space is 5D.

Nevertheless, the of 2/3 gained can be used to scale the savings when it comes to explore the conformational space associated with *m* tripeptides defining a flexible region along a protein backbone. For various plausible values of m, one obtains: (3/2)^5^ ~ 7.6 for *m* = 5, (3/2)^10^ ~ 57.7 for *m* = 10, (3/2)^15^ ~ 438 for *m* = 15, and (3/2)^20^ ~ 3325 for *m* = 20. Therefore, when *m* increases, an exponential reduction is the size of the search space to be explored is gained. This strategy is used in the companion paper [12]. Using depth *j* validity intervals with *j >* 1 can further reduce the search space as illustrated with the diminishing number of false positives (Fig. 8). Starting at depth 2 we have 18.12% fulfilling the constraint, then 16.57% for depth 3. There is no false positive in the planar dataset for depth 4 and onward.

**Figure 8:**
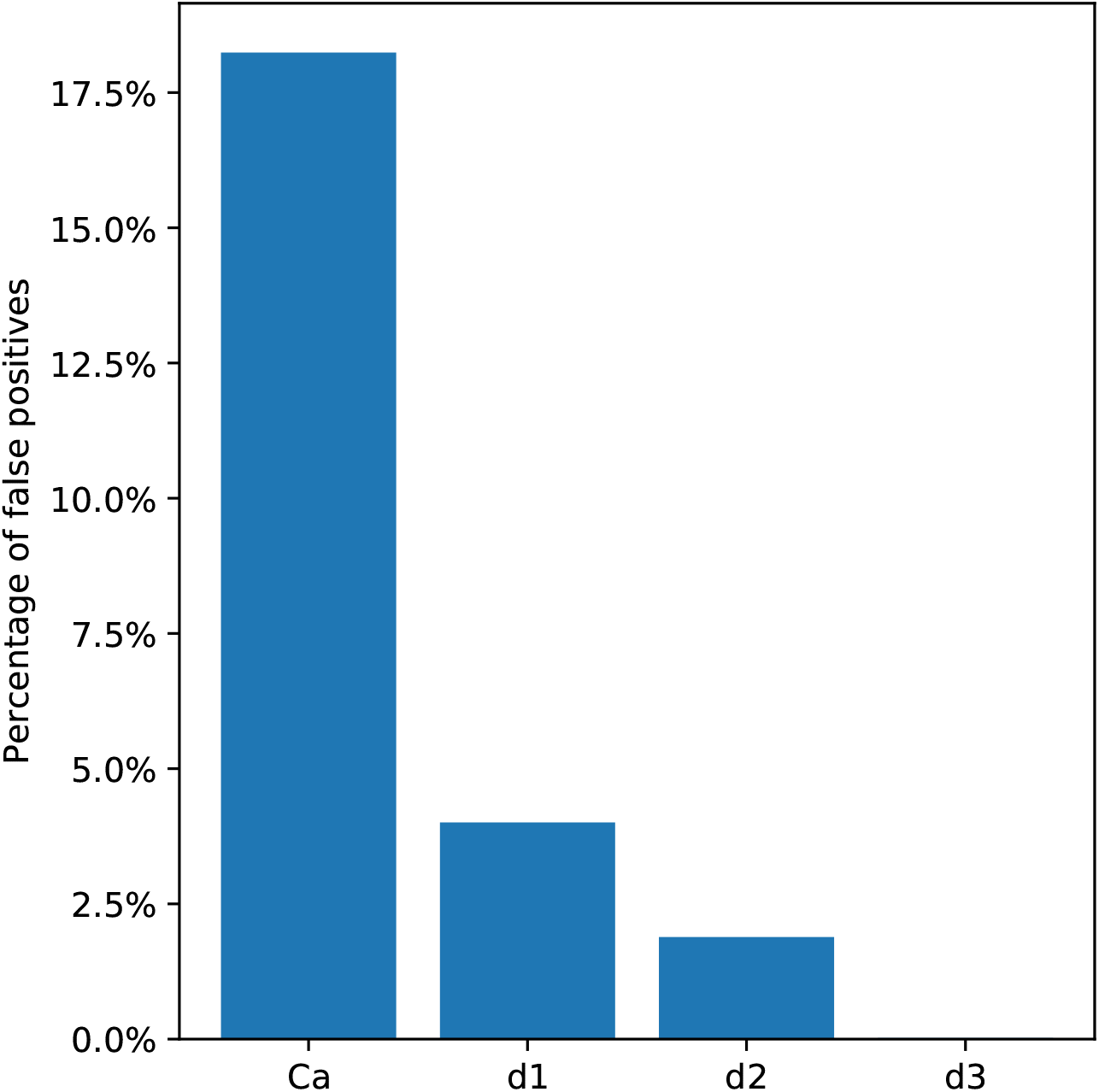
Proportion of false positives for *C_α_* valence and depth *j* validity intervals with *j* ∈ {1,2,3}. The proportion is defined as the number of false positives divided by the number instances when TLC yields no solution in the planar dataset (Sec. 5.1). The specific percentages are, *C_α_* valence constraint: 18.24%, depth 1 validity constraint: 4.01%, depth 2 validity constraint: 1.89%, depth 3 validity constraint: 0.02%.

## 6 Outlook

The compact nature of folded proteins makes the exploration of their conformational space especially challenging. One must indeed avoid steric clashes and optimize interactions, while dealing with coupled subproblems involving the backbone and side-chains. A critical need in this context are so-called movesets able to propose plausible (low energy) configurations given a starting pause. Such movesets are indeed a cornerstone in Monte Carlo based simulations at large.

The design of backbone movesets is a problem in itself, due to the necessity to handle loop closure constraints. The tripeptide loop closure provides an optimal solution to this problem, and therefore plays a crucial role to develop move sets. However, for loops involving several tripeptides, the question of combining solutions yielded by the individual tripeptides remains a challenging problem. Greedy approaches incrementally concatenating tripeptides have been developed, but these break the symmetry between the individual peptides, as the degree of freedom of those near the endpoints enjoy a finer sampling. On the way to processing all tripeptides in a sequence on an equal footing, this work studies (tight) necessary conditions on the first and last segment (bond) of a tripeptide, for TLC to yield solutions. As illustrated by our experiments, our conditions are rather tight, and yield an exponential saving in terms of the conformational space to be explored, when pooling several tripeptides. We leave the problem of improving the tightness of our constraints (notably using their iterated versions) as an open problem. Application-wise, the direct use of our constraints for the design of backbone movesets is presented in a companion paper.

## Acknowledgments

Théo Roncalli and Viraj Agashe are acknowledged for carefully re-reading the paper.

This work has been supported by the French government, through the 3IA Côte d’Azur Investments in the Future project managed by the National Research Agency (ANR) with the reference number ANR-19-P3IA-0002.

## Footnotes

1 Recall the triple product formula 〈*a, b* × *c*〉 = 〈(*a* × *b*), *c*〉.

