## Supporting Information for "Geometric constraints within tripeptides and the existence of tripeptide reconstructions"

### S1 Supporting information

#### S1.1 $C_\alpha$ valence constraint

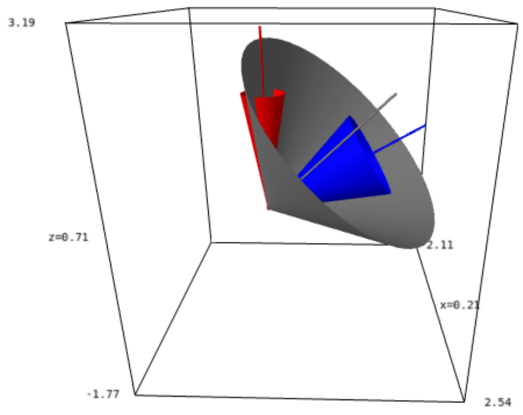

Figure S1: **Conservation of the valence angle $\theta_i = \angle\hat{\mathbf{r}}^{\sigma}_{\mathbf{i-1}}, \hat{\mathbf{r}}^{\tau}_{\mathbf{i}}$: illustration.** Angular values used: $\alpha_i = \angle\hat{\mathbf{Z}}_{\mathbf{i}}\hat{\mathbf{Z}}_{\mathbf{i-1}} = \mathbf{60}°$; blue cone of aperture $\xi_{i-1} = 20°$ representing the possible positions of the atom $N_i$, encoded using the vector $\hat{\mathbf{r}}^{\sigma}_{\mathbf{i-1}}$; red cone of aperture $\eta_i = 10°$ representing the possible positions of the atom $C_i$, encoded using the vector $\hat{\mathbf{r}}^{\tau}_{\mathbf{i}}$. Gray cone: a cone whose axis is on the blue cone – possible position for atom $N_i$, and with aperture $\theta_i$. The two intersections between the gray cone and the red one correspond to positions of $\hat{\mathbf{r}}^{\tau}_{\mathbf{i}}$ *i.e.* atom $C_i$, such that $\angle\hat{\mathbf{r}}^{\sigma}_{\mathbf{i-1}}, \hat{\mathbf{r}}^{\tau}_{\mathbf{i}} = \theta_i$.

#### S1.2 Viète's law and limit values for the angles $\sigma_{i-1;-}$ and $\sigma_{i-1;+}$

We establish the extreme values for $\langle \hat{\mathbf{r}}^{\sigma}_{\mathbf{i-1}}, \hat{\mathbf{Z}}_{\mathbf{i}} \rangle$, that is

$$\langle \hat{\mathbf{r}}^{\sigma}_{\mathbf{i-1}}, \hat{\mathbf{Z}}_{\mathbf{i}} \rangle = \cos(\theta_{\mathbf{i}} \pm \eta_{\mathbf{i}}).$$

We use spherical trigonometry (Fig. S2)(A). Using Viète's law of cosines for the spherical triangle $ABC$, we get

$$\cos x = \cos\theta_i \cos\eta_i + \sin\theta_i \sin\eta_i \cos\gamma. \tag{44}$$

The extreme values for $\gamma$ are 0 and $\pi$, and the corresponding extreme values $\cos(\theta_i - \eta_i)$ and $\cos(\theta_i + \eta_i)$ respectively (Fig. S2)(B).

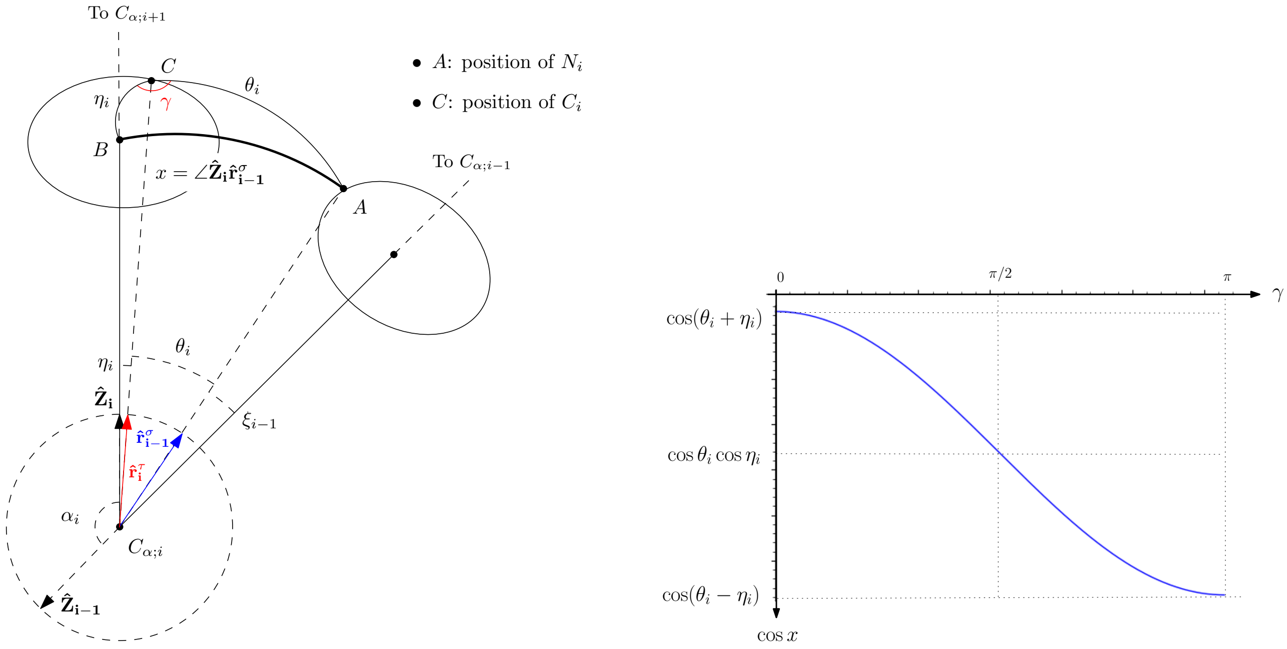

Figure S2: **Conservation of the angle $\theta_i$. (A)** Viète formula of cosines. Note that the triangle $ABC$ is defined by great circle arcs on $S^2$. **(B)** Variation of $\cos x$ – curve plotted assuming with $\theta_i = 116°$ and $\eta_i = 10°$.

#### S1.3 Signatures

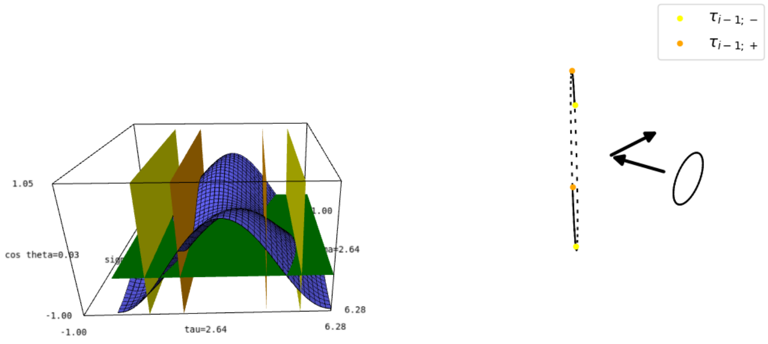

PNZZ

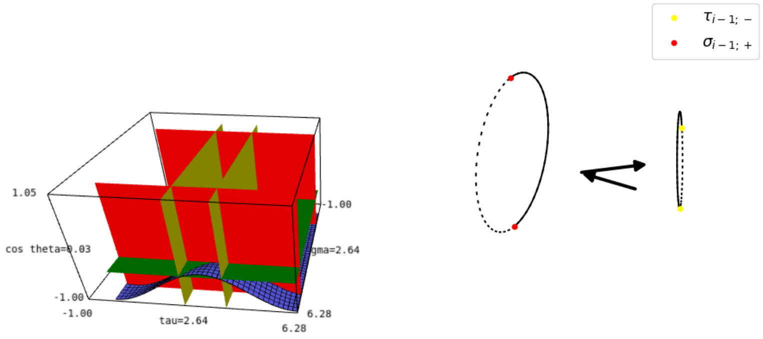

PZZN

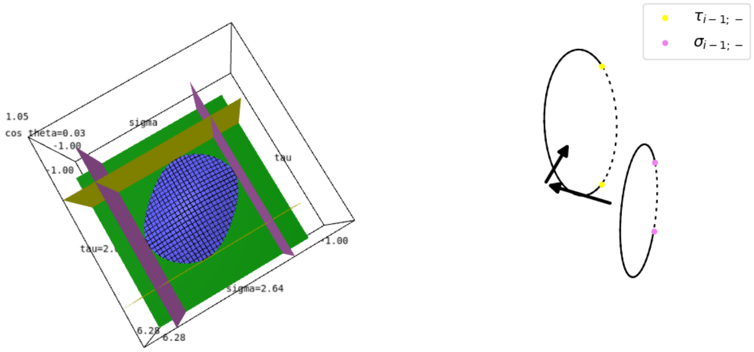

ZNZN

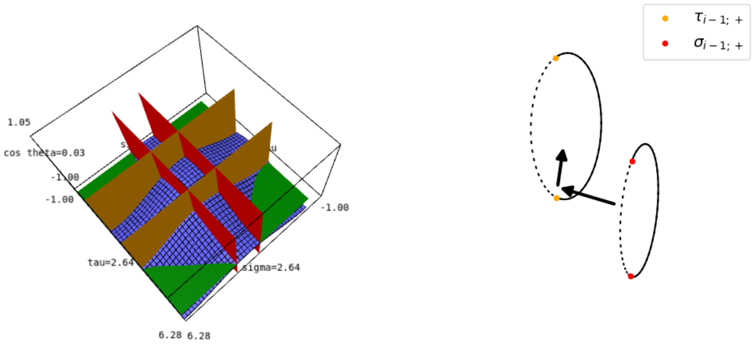

PZPZ

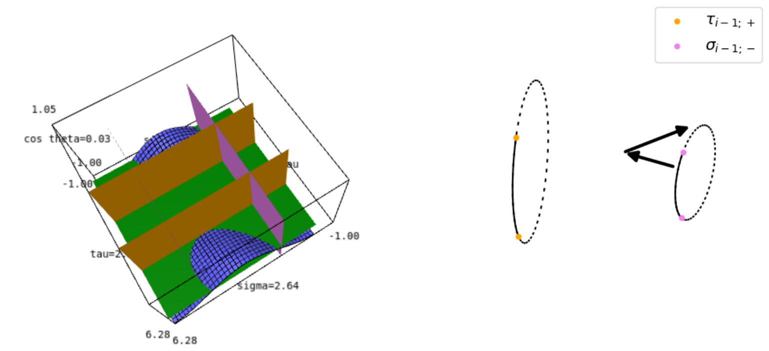

ZNPZ

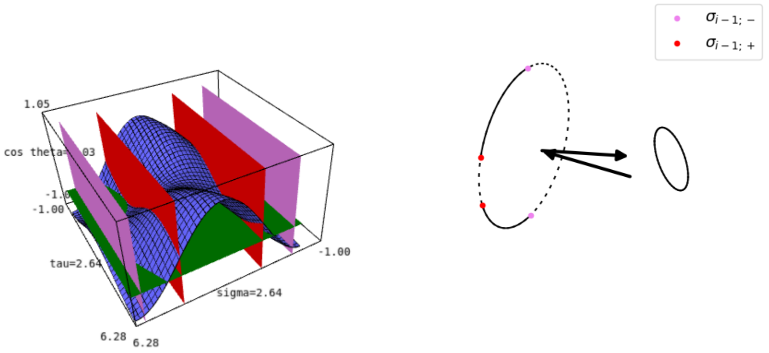

ZZPN

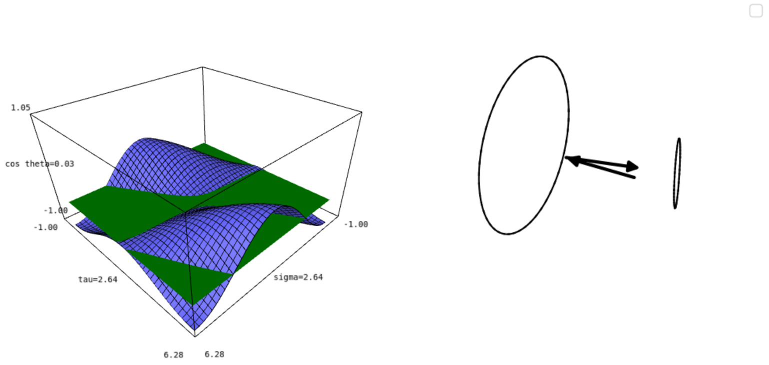

PNPN

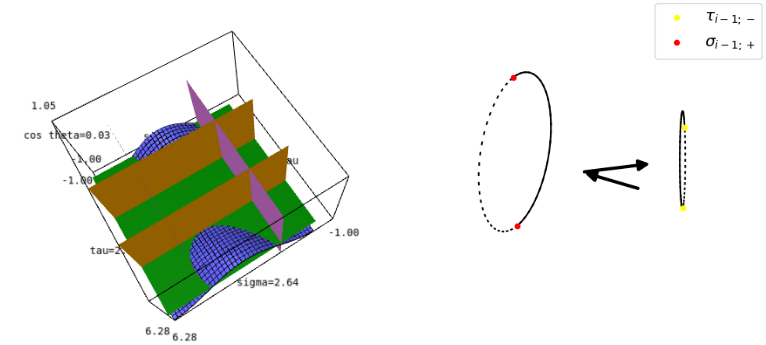

ZNPZ

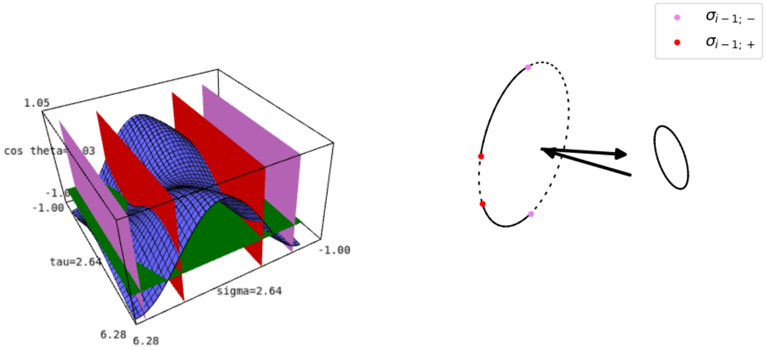

ZZPN

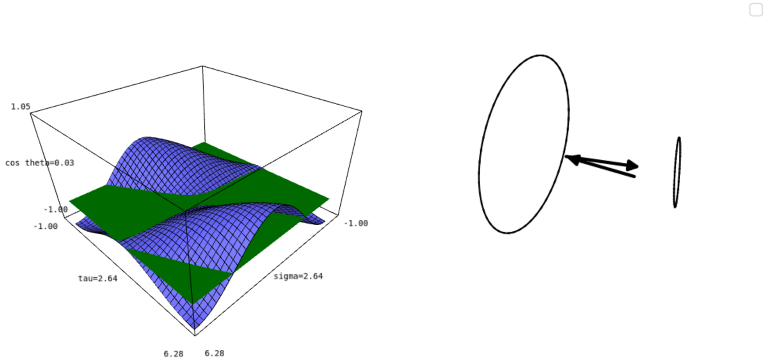

PNPN

Figure S3: $\sigma_{i-1}$ $\tau_i$ **dot surfaces and validity intervals for the dataset of random TLC instances.** **(A)** The 7 signatures (Def. 7) in terms of extreme angles for the data set of random TLC instances. In all cases, the green plane corresponds to $\cos\theta_i = \cos 111.6°$. A signature reads as follows: N:negative ie dot product $< -1$; Z: zero ie dot product $\in [-1, 1]$; P: positive ie dot product $> 1$. The vertical planes correspond to extreme angles. The color code is the same as that of **(B)**. **(B)** Validity intervals as arc on the same instances. Black bold line for valid intervals, dashed line otherwise. The colored bullets indicate the corresponding extreme angle as defined in the legend.

#### S1.4 Functions $\sigma^*_{i-1}(\tau_i)$ and $\sigma^{**}_{i-1}(\tau_i)$

We present the derivation for functions $\sigma^*_{i-1}(\tau_i)$ and $\sigma^{**}_{i-1}(\tau_i)$ from Eq. (40). The case of $\tau^*_i(\sigma_{i-1})$ and $\tau^{**}_i(\sigma_{i-1})$ is similar, and thus omitted.

##### S1.4.1 Functions

The dot product equations for both $\sigma_{i-i}$ and $\tau_i$ are of the general form:

$$K^{(1)}_{\sigma_{i-1}} \cos\sigma_{i-1} + K^{(2)}_{\sigma_{i-1}} \sin\sigma_{i-1} + K^{(3)}_{\sigma_{i-1}} = 0 \tag{45}$$

We divide by $\sqrt{{K^{(1)}_{\sigma_{i-1}}}^2 + {K^{(2)}_{\sigma_{i-1}}}^2}$ and define two angles:

$$\begin{cases} B = \operatorname{atan2}\left(K^{(2)}_{\sigma_{i-1}}, K^{(1)}_{\sigma_{i-1}}\right) \\ C = \arccos\left(\frac{-K^{(3)}_{\sigma_{i-1}}}{\sqrt{{K^{(1)}_{\sigma_{i-1}}}^2 + {K^{(2)}_{\sigma_{i-1}}}^2}}\right) \end{cases} \tag{46}$$

Note that $C$ is only well-defined when $|K^{(3)}_{\sigma_{i-1}}| \leq \sqrt{{K^{(1)}_{\sigma_{i-1}}}^2 + {K^{(2)}_{\sigma_{i-1}}}^2}$. Using the trigonometric identity $\cos(a-b) = \cos a \cos b + \sin a \sin b$, we have:

$$\cos(\sigma_{i-1} - B) = \cos C \tag{47}$$

The general solution of this equation is $\sigma_{i-1} = 2n\pi + B \pm C$ for $n \in \mathbb{Z}_\sigma$. Since we are considering the solutions modulo $2\pi$, as reported in Eq. (40).

##### S1.4.2 Delineating values for atan2

**Function** atan2. Using the previous equations, the solutions of the dot product equations are given by:

$$\begin{cases} \sigma^*_{i-1}(\tau_i) = \operatorname{atan2}\left(K^{(2)}_{\sigma_{i-1}}, K^{(1)}_{\sigma_{i-1}}\right) + \arccos\left(\frac{-K^{(3)}_{\sigma_{i-1}}}{\sqrt{{K^{(1)}_{\sigma_{i-1}}}^2 + {K^{(2)}_{\sigma_{i-1}}}^2}}\right) \\ \sigma^{**}_{i-1}(\tau_i) = \operatorname{atan2}\left(K^{(2)}_{\sigma_{i-1}}, K^{(1)}_{\sigma_{i-1}}\right) - \arccos\left(\frac{-K^{(3)}_{\sigma_{i-1}}}{\sqrt{{K^{(1)}_{\sigma_{i-1}}}^2 + {K^{(2)}_{\sigma_{i-1}}}^2}}\right) \end{cases} \tag{48}$$

For ease of plotting and calculations, the atan2 function is expressed in terms of the standard arctan function as follows:

$$\operatorname{atan2}(y,x) = \begin{cases} \arctan(\frac{y}{x}) & x > 0 \\ \arctan(\frac{y}{x}) + \pi & x < 0, y \geq 0 \\ \arctan(\frac{y}{x}) - \pi & x < 0, y < 0 \\ \frac{\pi}{2} & x = 0, y \geq 0 \\ -\frac{\pi}{2} & x = 0, y < 0 \\ \text{undefined} & x = 0, y = 0 \end{cases} \tag{49}$$

**Application to our case.** Recalling the definitions of $K^{(1)}_{\sigma_{i-1}}, K^{(2)}_{\sigma_{i-1}}$:

$$\begin{cases} K^{(1)}_{\sigma_{i-1}} & = \cos\tau_i \sin\xi_{i-1} \sin\eta_i \cos\alpha_i + \sin\alpha_i \cos\eta_i \sin\xi_{i-1} \\ K^{(2)}_{\sigma_{i-1}} & = \sin\xi_{i-1} \sin\eta_i \sin\tau_i \end{cases} \tag{50}$$

To apply Eq. (49) in our case, we discuss the signs of $K^{(1)}_{\sigma_{i-1}}$ and $K^{(2)}_{\sigma_{i-1}}$, noting that $\xi_{i-1}$ and $\eta_i \in (0, \pi)$.

As will be clear shortly, we also denote $Z_\sigma = \arccos\left(-\frac{\tan\alpha_i}{\tan\eta_i}\right)$.

**Condition for $K^{(2)}_{\sigma_{i-1}} < 0$.** Plainly, $K^{(2)}_{\sigma_{i-1}} < 0 \implies \tau_i \in (\pi, 2\pi)$.

**Condition for $K^{(1)}_{\sigma_{i-1}} > 0$.** We distinguish two cases: • When $\cos\alpha_i > 0$, since $\sin\xi_{i-1} > 0$:

$$K^{(1)}_{\sigma_{i-1}} > 0 \Leftrightarrow \cos\tau_i \geq -\frac{\sin\alpha_i \cos\eta_i}{\cos\alpha_i \sin\eta_i} = -\frac{\tan\alpha_i}{\tan\eta_i} \Leftrightarrow \tau_i \in (0, Z_\sigma) \cup (2\pi - Z_\sigma, 2\pi) \tag{51}$$

• When $\cos\alpha_i < 0$, since $\sin\xi_{i-1} > 0$:

$$K^{(1)}_{\sigma_{i-1}} > 0 \Leftrightarrow \cos\tau_i \leq -\frac{\sin\alpha_i \cos\eta_i}{\cos\alpha_i \sin\eta_i} = -\frac{\tan\alpha_i}{\tan\eta_i} \Leftrightarrow \tau_i \in (Z_\sigma, 2\pi - Z_\sigma) \tag{52}$$

**Final expression.** Therefore we may express $\operatorname{atan2}\left(K^{(2)}_{\sigma_{i-1}}, K^{(1)}_{\sigma_{i-1}}\right)$ in terms of the intervals of $\tau_i$ as follows.
• When $\cos\alpha_i > 0$:

$$\operatorname{atan2}\left(K^{(2)}_{\sigma_{i-1}}, K^{(1)}_{\sigma_{i-1}}\right) = \begin{cases} \arctan\left(\frac{K^{(2)}_{\sigma_{i-1}}}{K^{(1)}_{\sigma_{i-1}}}\right) & \tau_i \in [0, Z_\sigma) \cup (2\pi - Z_\sigma, 2\pi] \\ \arctan\left(\frac{K^{(2)}_{\sigma_{i-1}}}{K^{(1)}_{\sigma_{i-1}}}\right) + \pi & \tau_i \in (Z_\sigma, \pi] \\ \arctan\left(\frac{K^{(2)}_{\sigma_{i-1}}}{K^{(1)}_{\sigma_{i-1}}}\right) - \pi & \tau_i \in (\pi, 2\pi - Z_\sigma) \\ \frac{\pi}{2} & \tau_i = Z_\sigma \\ -\frac{\pi}{2} & \tau_i = 2\pi - Z_\sigma \end{cases}$$

• When $\cos\alpha_i < 0$:

$$\operatorname{atan2}\left(K^{(2)}_{\sigma_{i-1}}, K^{(1)}_{\sigma_{i-1}}\right) = \begin{cases} \arctan\left(\frac{K^{(2)}_{\sigma_{i-1}}}{K^{(1)}_{\sigma_{i-1}}}\right) & \tau_i \in (Z_\sigma, 2\pi - Z_\sigma) \\ \arctan\left(\frac{K^{(2)}_{\sigma_{i-1}}}{K^{(1)}_{\sigma_{i-1}}}\right) + \pi & \tau_i \in [0, Z_\sigma) \\ \arctan\left(\frac{K^{(2)}_{\sigma_{i-1}}}{K^{(1)}_{\sigma_{i-1}}}\right) - \pi & \tau_i \in (Z_\sigma, 2\pi) \\ \frac{\pi}{2} & \tau_i = Z_\sigma \\ -\frac{\pi}{2} & \tau_i = 2\pi - Z_\sigma \end{cases}$$

We can similarly obtain the corresponding expressions for the other angle. Let $Z_\tau = \arccos\left(-\frac{\tan\alpha_i}{\tan\xi_{i-1}}\right)$. The intervals are then given by:
• When $\cos\alpha_i > 0$:

$$\operatorname{atan2}\left(K^{(2)}_{\tau_i}, K^{(1)}_{\tau_i}\right) = \begin{cases} \arctan\left(\frac{K^{(2)}_{\tau_i}}{K^{(1)}_{\tau_i}}\right) & \sigma_{i-1} \in [0, Z_\tau) \cup (2\pi - Z_\tau, 2\pi] \\ \arctan\left(\frac{K^{(2)}_{\tau_i}}{K^{(1)}_{\tau_i}}\right) + \pi & \sigma_{i-1} \in (Z_\tau, \pi] \\ \arctan\left(\frac{K^{(2)}_{\tau_i}}{K^{(1)}_{\tau_i}}\right) - \pi & \sigma_{i-1} \in (\pi, 2\pi - Z_\tau) \\ \frac{\pi}{2} & \sigma_{i-1} = Z_\tau \\ -\frac{\pi}{2} & \sigma_{i-1} = 2\pi - Z_\tau \end{cases}$$

- When $\cos\alpha_i < 0$:

$$\operatorname{atan2}\left(K^{(2)}_{\tau_i}, K^{(1)}_{\tau_i}\right) = \begin{cases} \arctan\left(\frac{K^{(2)}_{\tau_i}}{K^{(1)}_{\tau_i}}\right) & \sigma_{i-1} \in (Z_\tau, 2\pi - Z_\tau) \\ \arctan\left(\frac{K^{(2)}_{\tau_i}}{K^{(1)}_{\tau_i}}\right) + \pi & \sigma_{i-1} \in [0, Z_\tau) \\ \arctan\left(\frac{K^{(2)}_{\tau_i}}{K^{(1)}_{\tau_i}}\right) - \pi & \sigma_{i-1} \in (Z_\tau, 2\pi) \\ \frac{\pi}{2} & \sigma_{i-1} = Z_\tau \\ -\frac{\pi}{2} & \sigma_{i-1} = 2\pi - Z_\tau \end{cases}$$

#### S1.5 Plots

Consider the functions $\sigma_{i-1}(\tau_i), \tau_i(\sigma_{i-1})$, and $\tau_i^*(\sigma_{i-1}), \tau_i^{**}(\sigma_{i-1})$. We plot their graph and that of their derivatives for instances of the seven signatures (Def. 7) observed. Conventions on the plots are as follows:

- Left column: the blue and red curves represent the two functions $\sigma^*_{i-1}(\tau_i)$ and $\sigma^{**}_{i-1}(\tau_i)$ respectively. The orange and cyan curves represent their derivatives respectively.
- Right column: the blue and red curves represent the two functions $\tau_i^*(\sigma_{i-1})$ and $\tau_i^{**}(\sigma_{i-1})$ respectively. The orange and cyan curves represent their derivatives respectively.
- The purple bar denotes the extent of the initial validity interval in $[0, \pi]$ and the yellow bar denotes the extent of the initial validity intervals in $[\pi, 2\pi)$.

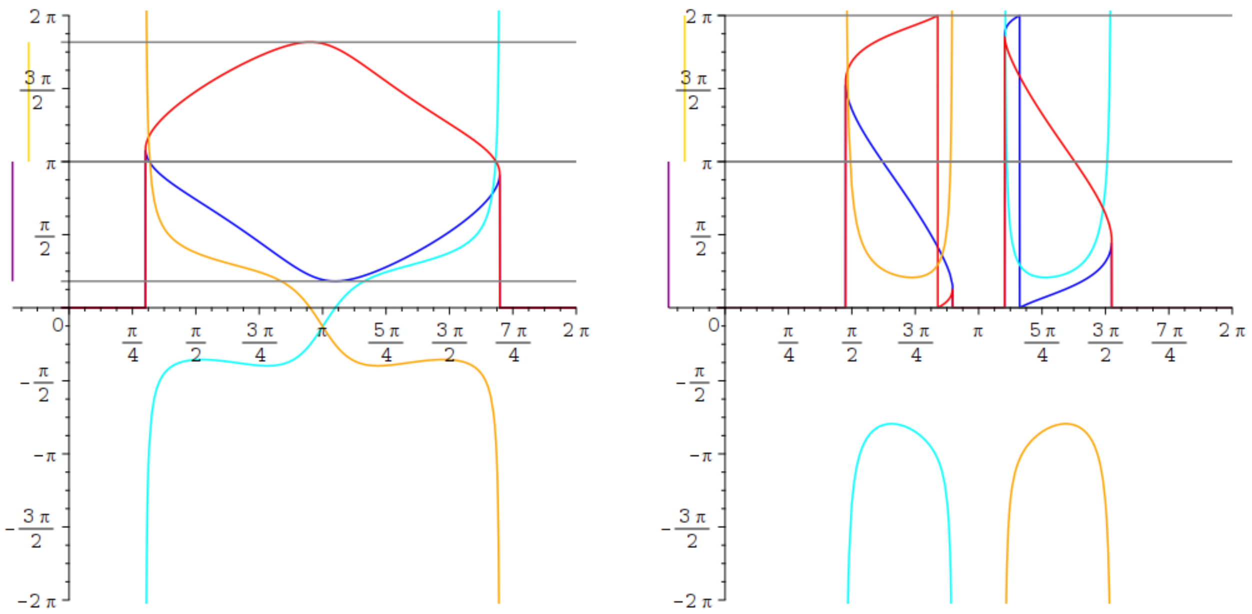

$\sigma_{i-1}(\tau_i)$ $\tau_i(\sigma_{i-1})$

Figure S4: Signature PNZZ

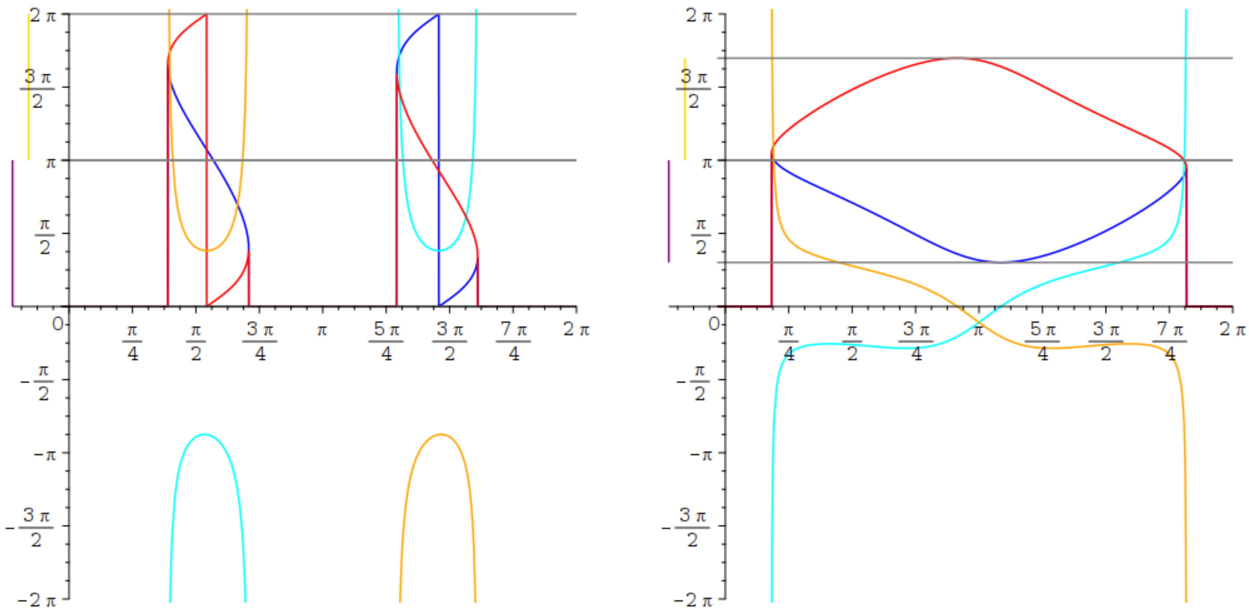

$\sigma_{i-1}(\tau_i)$ $\tau_i(\sigma_{i-1})$

Figure S5: Signature ZNZN

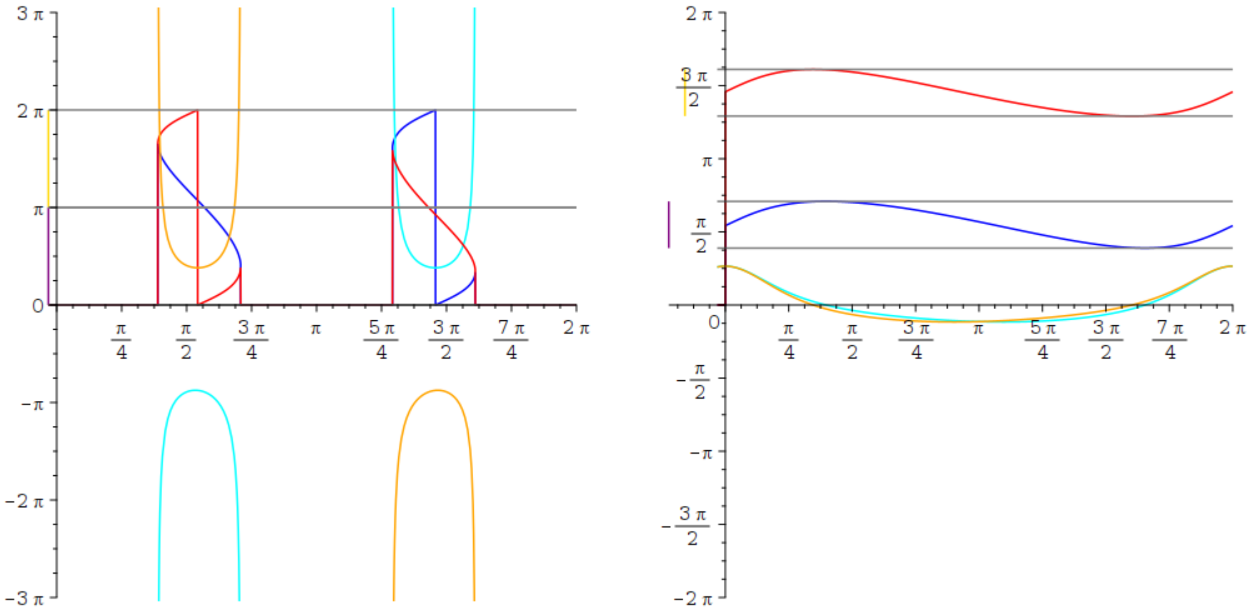

$\sigma_{i-1}(\tau_i)$ $\tau_i(\sigma_{i-1})$

Figure S6: Signature ZZPN

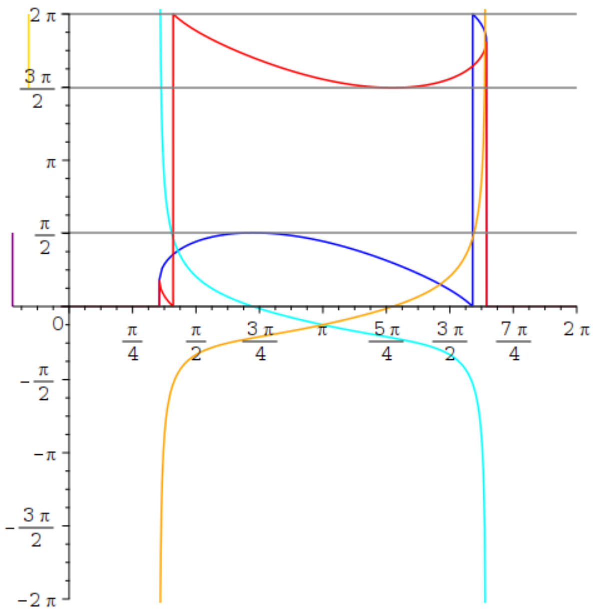

$\sigma_{i-1}(\tau_i)$ $\tau_i(\sigma_{i-1})$

Figure S7: Signature ZNPZ

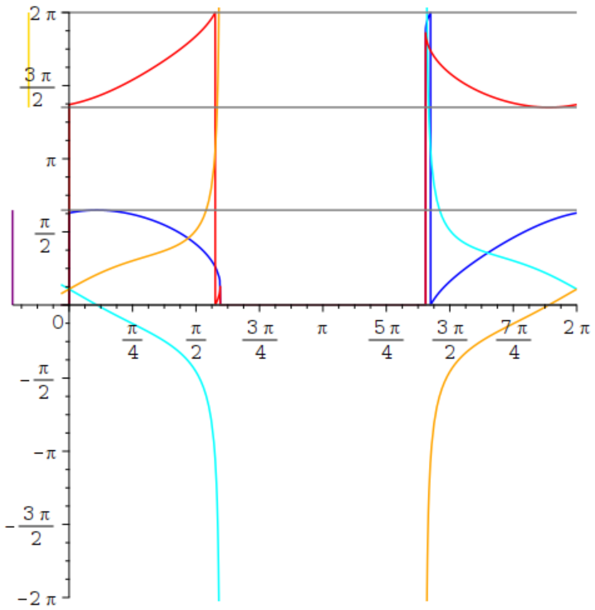

$\sigma_{i-1}(\tau_i)$ $\tau_i(\sigma_{i-1})$

Figure S8: Signature PZPZ

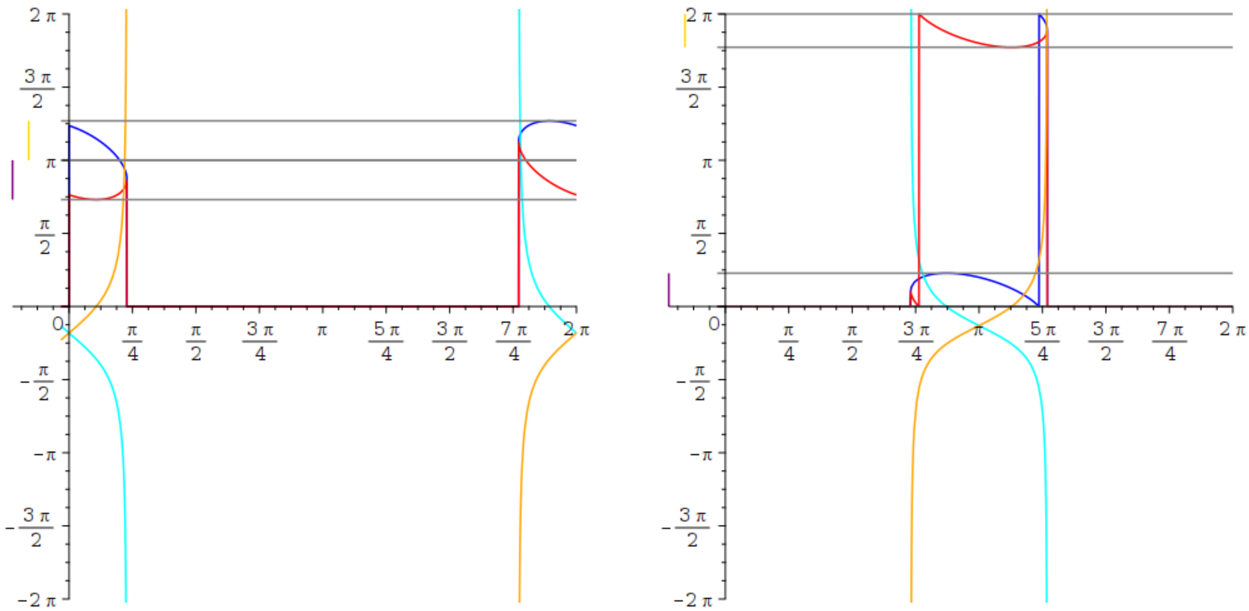

$\sigma_{i-1}(\tau_i)$ $\tau_i(\sigma_{i-1})$

Figure S9: Signature PZZN

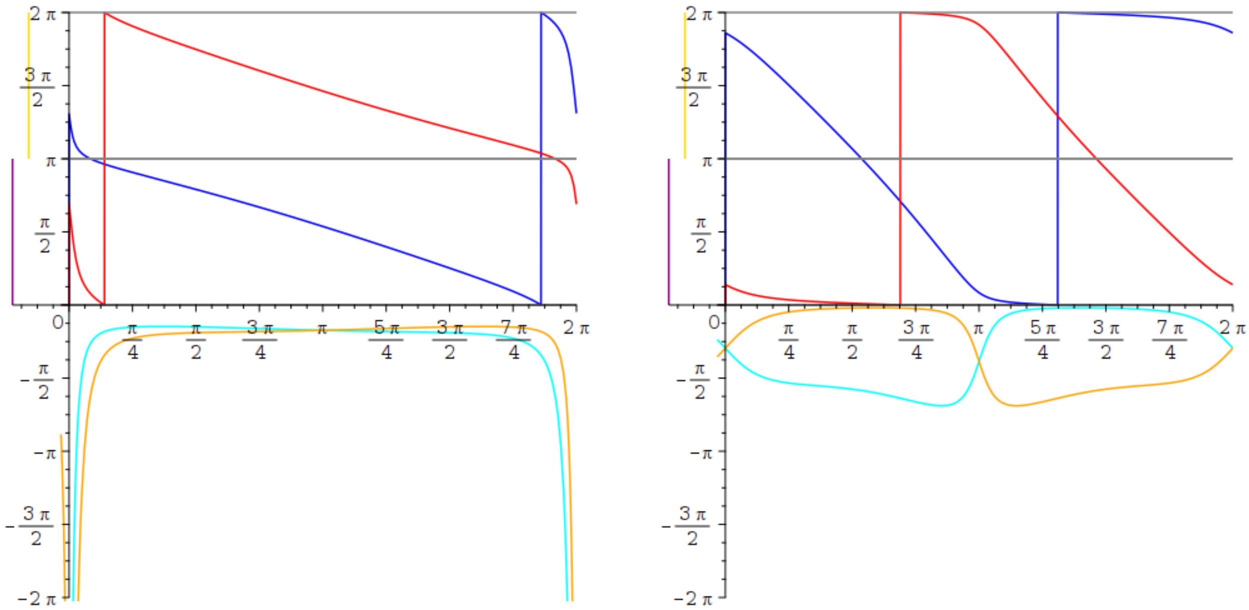

$\sigma_{i-1}(\tau_i)$ $\tau_i(\sigma_{i-1})$

Figure S10: Signature PNPN

**6 Outlook** **13**

**7 Artwork** **14**

**S1 Supporting information** **21**
- S1.1 $C_\alpha$ valence constraint . . . 21
- S1.2 Viète's law and limit values for the angles $\sigma_{i-1;-}$ and $\sigma_{i-1;+}$ . . . 22
- S1.3 Signatures . . . 22
- S1.4 Functions $\sigma^*_{i-1}(\tau_i)$ and $\sigma^{**}_{i-1}(\tau_i)$ . . . 26
  - S1.4.1 Functions . . . 26
  - S1.4.2 Delineating values for atan2 . . . 26
- S1.5 Plots . . . 28
